# Introduced chemokine gradients guide transplanted and regenerated retinal neurons toward their natural position in the retina

**DOI:** 10.1101/2022.09.29.510158

**Authors:** Jonathan R Soucy, Levi Todd, Emil Kriukov, Monichan Phay, Thomas A Reh, Petr Baranov

**Author notes:** Corresponding authors: Jonathan R Soucy; Petr Baranov.

## Abstract

Ongoing cell replacement studies and clinical trials have demonstrated the need to control donor and newborn cell behavior within their target tissue. Here we present a methodology to guide stem cell-derived and endogenously regenerated neurons by engineering the microenvironment. Being an “approachable part of the brain,” the eye provides a unique opportunity to study donor neuron fate, migration, and integration within the central nervous system. Glaucoma and other optic neuropathies lead to the permanent loss of retinal ganglion cells (RGCs) – the neurons in the retina that transfer all visual information from the eye to the brain. Cell transplantation and transdifferentiation strategies have been proposed to restore RGCs, and one of the significant barriers to successful RGC integration into the existing retinal circuitry is cell migration towards their natural position in the retina. Here we describe a framework for identifying, selecting, and applying chemokines to direct cell migration in vivo within the retina. We have performed an in silico analysis of the single-cell transcriptome of the developing human retina and identified six receptor-ligand candidates to guide stem cell-derived or newborn neurons. The lead candidates were then tested in functional in vitro assays for their ability to guide stem cell-derived RGCs. For the in vivo studies, donor and newborn neurons were differentiated in human and mouse retinal organoids or endogenously reprogrammed with proneuronal transcription factors, respectively. An exogenous stromal cell-derived factor-1 (SDF1) gradient led to a 2.7-fold increase in donor RGC migration into the ganglion cell layer and a 3.3-fold increase in the displacement of newborn RGCs out of the inner nuclear layer. Furthermore, by altering the migratory profile of donor RGCs toward multipolar migration, overall migration was improved in mature retinal tissues. Together, these results highlight the ability and importance of engineering the tissue microenvironment and the individual cells for research and clinical applications in gene and cell therapies.

**Graphical Abstract:** 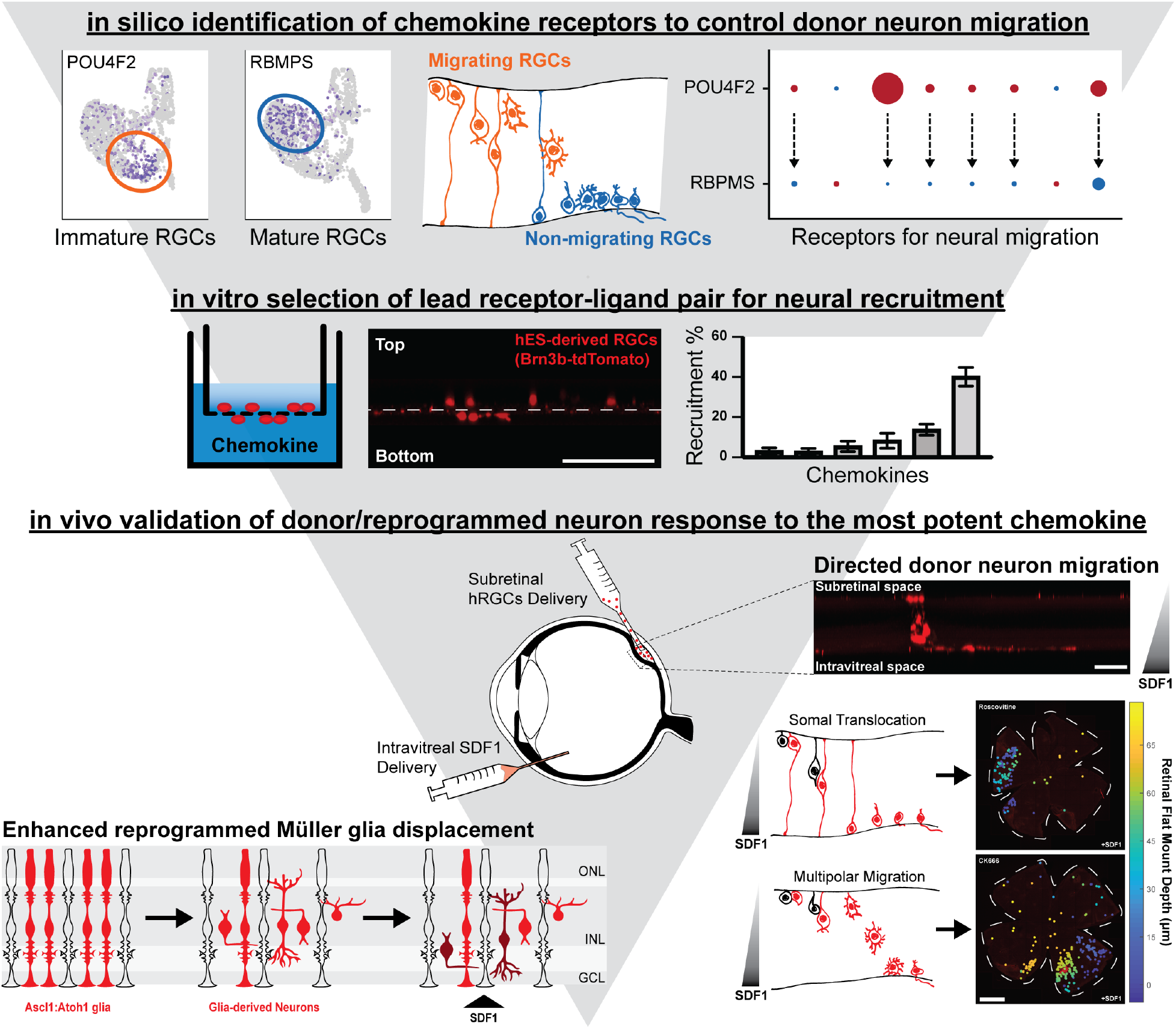

In brief, the “in silico – in vitro – in vivo” funnel holds significant potential for identifying targets to control cellular processes in research and clinical applications. In this report, Soucy et al. describes a framework for identifying, selecting, and applying chemokines to direct retinal ganglion cell migration in vivo within the adult mouse retina.

---

**A**pproximately 3.5% of the world’s population over 40 years old has glaucoma, the most common optic neuropathy.^1^ By 2040, the global prevalence of glaucoma will exceed 110 million people as the population ages,^2^ making it a high-priority target for therapy development. Currently, there is no treatment to restore lost visual function. Although several neuroprotective approaches focused on preserving existing cells and their axons are being explored;^3,4^ this strategy alone will not restore vision that is already lost due to cell death. The mammalian retina has limited capacity to regenerate; thus, retinal neuron death leads to irreversible vision loss.^5^ RGC replacement is needed to recover sight loss to glaucoma.

Retinal ganglion cell (RGC) replacement remains an unsolved challenge in regenerative ophthalmology. Success would help bring vision back to millions of advanced-state glaucoma patients. Recent advancements enable transplantation of primary rodent RGCs,^6–8^ differentiation of RGCs from human pluripotent stem cells,^9,10^ reprogramming Müller glia to RGCs,^11^ and functional axon regeneration to the brain suggest that transplantation-mediated vision repair may be feasible.^12^

We were among the first to demonstrate the robust survival of induced pluripotent stem cell (iPSC)-derived RGCs following intravitreal (IVT) transplantation into healthy and damaged retinas.^13^ These studies became possible due to robust RGC differentiation and isolation from iPSC cultures established in our lab.^14–16^ Despite those successes, the survival rate for individual RGCs remains low, and most donor neurons remain above the internal limiting membrane (ILM) that defines the neural retinal border with the vitreous cavity without integrating.^17^ This challenge is not unique to RGC replacement. Poor structural integration is a significant obstacle to neuron transplantation (e.g., photoreceptors, dopaminergic, and motor neurons) and transdifferentiation (e.g., glia to cerebral and retinal neurons).^11,18^ The recent discovery of techniques for non-surgical ILM disruption of the host removes the existing anatomical barrier;^19^ however, integration critically depends on the modulation of the host’s adult retinal environment, and ILM disruption may not be appropriate in a clinical setting.^20^ We hypothesize that early guided migration can significantly improve the structural and functional integration of donor and newborn RGCs.

Neurogenesis in the mammalian retina completes shortly after birth.^21,22^ Thus, transplantation of stem cells and their progeny relies on the recapitulation of the development and/or regenerative pathways. During development, RGCs, like most early-born neurons, migrate via somal translocation (ST), but confocal traces in zebrafish demonstrate that RGCs migrate through multipolar (MP) migration if ST is inhibited.^23^ MP migration does not rely on the extension and attachment of neural processes to reach their final location and is the preferred migratory mode for late-born neurons that navigate through developed tissues.^24,25^ It is unknown if RGCs are capable of MP migration in mammals, and the exact mechanism by which stem cell-derived donor RGCs migrate within the mature retina remains unknown. It is also not clear if newborn and stem cell-derived RGCs can respond to chemokines, previously identified in the studies of retinal and cerebral development and on neuron migration in the brain out of the subventricular zone during postnatal regeneration.^26^

Here we describe a framework to identify, select, and apply chemokines to direct cell migration in vivo within the retina. We performed an in silico analysis of the single-cell transcriptome of developing human retinas and identified six receptor-ligand candidates to guide stem cell-derived or newborn neurons. The lead candidates were then tested in the functional in vitro assays for their ability to guide stem cell-derived RGCs, with stromal cell-derived factor-1 (SDF1) identified as the most potent chemokine for RGC recruitment. For this and other experiments, we differentiated RGCs from mouse and human stem cells using retinal organoid cultures or stimulated glial reprogramming to neurons with proneuronal transcription factors.^11,27,28^ We then transplanted these stem cell-derived RGCs subretinally and delivered recombinant SDF1 protein intravitreally to establish a chemokine gradient across the retina. Using a quantitative approach to transplantation, we confirmed that donor cell behavior is controllable by modulating the tissue microenvironment. Furthermore, we demonstrate that an SDF1 gradient across the host retina enhances the structural integration of mouse and human stem cell-derived donor RGCs via MP migration. Lastly, we demonstrate that IVT delivery of SDF1 increases the displacement of newborn RGCs out of the inner nuclear layer and towards their natural connecting points in the retina. Altogether, this is the first demonstration of the universal nature and applicability of neurokine-directed controlled migration of donor stem cell-derived and endogenously regenerated neurons. Moreover, the established workflow to identify microenvironment modifiers can be ported to control other aspects of donor neuron behavior.

## Results and Discussion

### Donor stem cell-derived RGCs fail to spontaneously migrate into the retina

We and others have demonstrated the feasibility of cell replacement therapy with RGCs isolated from the developing retina and stem cell-derived RGCs. Our grafts survived in healthy and damaged retinas following xeno- and allo-transplantation and sent projections into the optic nerve.^29^ As cell survival does not equal functional integration, better structural integration is needed to achieve vision restoration. While >60% of recipients have donor RGCs at two weeks to one year in our syngeneic transplantation study,^29^ the proportion of surviving cells typically remains low (<5%), and even fewer cells migrate completely into the ganglion cell layer (<1%). Recently, Zhang et al. showed that stem cell-derived human RGCs cultured on the neuroretina explant could not migrate through the internal limiting membrane (ILM) to graph into the GCL, and disruption of this barrier resulted in increased structural integration.^48^ However, while it is possible to degrade or surgically remove the ILM, these procedures pose a significant risk of damaging the retina.^20^ Moreover, the ILM has been shown to play an essential role in the proper lamination of the retina.^49^ Therefore, we sought to explore the subretinal (SR) delivery as an alternative approach for transplanting donor neurons while leaving the ILM completely intact.

SR delivery has proven an effective strategy for gene, photoreceptor, and retinal pigment epithelium delivery. However, unlike these transplantation paradigms, where the cells are delivered directly to their final position, RGCs must migrate > 200 μm into the GCL. While our previous work suggests that the SR space supports donor RGC survival, we previously did not observe any migration towards the GCL using this approach.^29^ During development, RGCs are born on the apical surface and migrate towards the basal side into what will become the GCL, but we do not yet know if this is due to an intrinsic RGC capacity or a response to the developmental retinal microenvironment. To investigate this phenomenon, we delivered mouse stem cell-derived RGCs using an intravitreal (IVT) and SR approach to study the positions of the donor cells within the neural retina. Two weeks post-transplantation, mice were euthanized, and retinas were stained and mounted to access donor RGC distribution (Fig. 1A). We observed no spontaneous migration through the ILM for RGCs delivered IVT (Fig. 1B) but demonstrated for the first-time limited migration through the neural retina for RGCs delivered subretinally (Fig. 1C). Despite having a limited capacity for spontaneous migration following SR cell delivery, and very few RGCs integrating into the GCL, these results suggest that the SR delivery route is a viable alternative to IVT delivery so that RGCs can circumvent the ILM. Moreover, because spontaneous migration only occurred by mimicking development with our SR delivery approach, we hypothesize that neuron migration in the retina must be driven by a response to the developmental microenvironment.

**Fig. 1.**
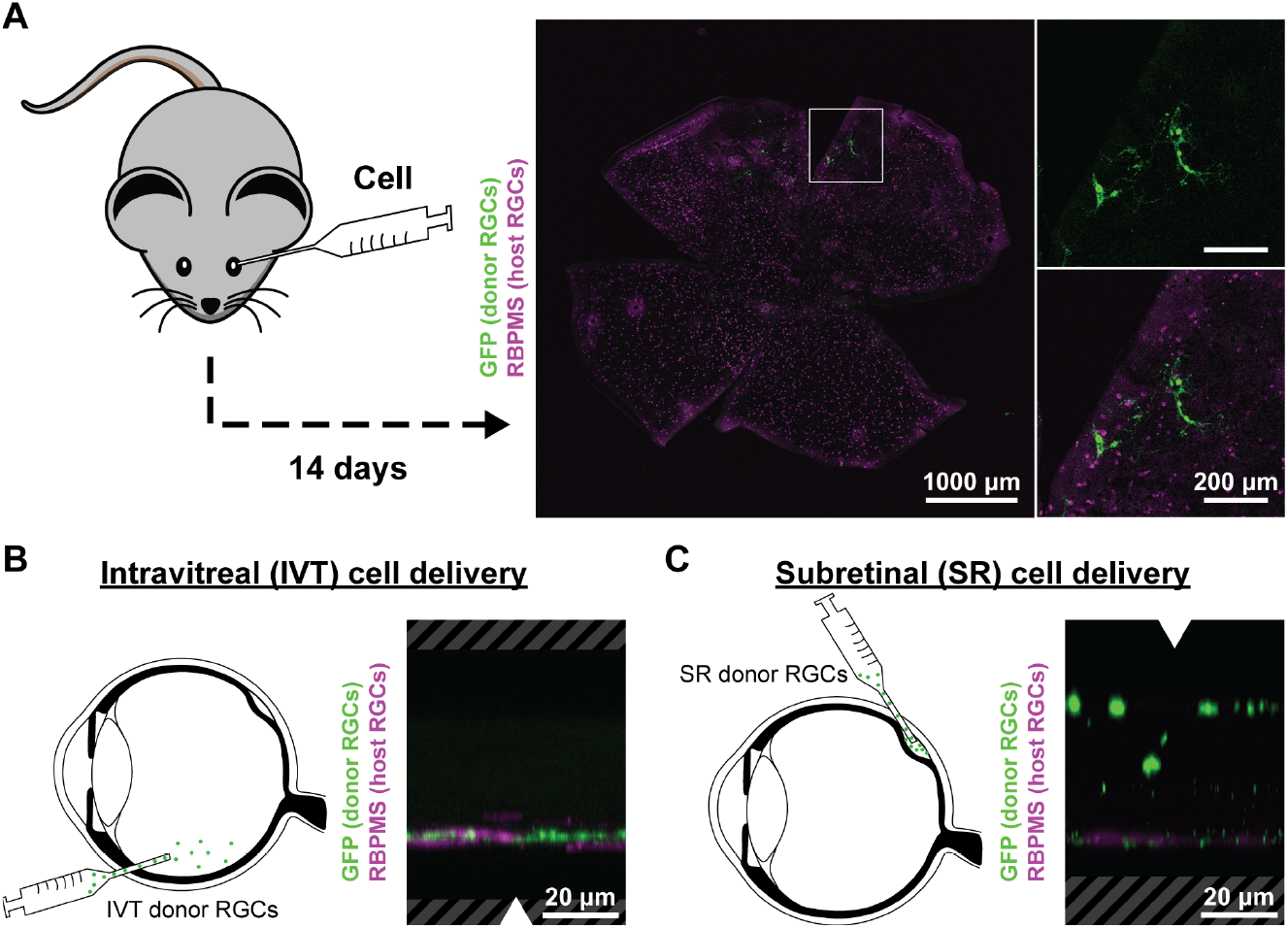
Spontaneous RGC migration towards the GCL following subretinal delivery in mice. **(A)** Representative immunofluorescent image of a mouse retinal flat mount (max intensity projection) stained for host (mThy1.2-GFP+) and donor (RBMPS+) RGCs 14 days post-transplantation. **(B)** Representative orthographic projection of a retinal flat mount following IVT delivery and **(C)** SR delivery of donor RGCs.

### Identifying migration cues through single-cell RNA sequencing analysis of the developing and adult human retina

To investigate RGC migration in the developing retina, we sourced available human fetal and adult retina single-cell RNA sequencing data to explore which pro-migratory signals are upregulated during RGC development.^32,33^ There are two major neuron migration patterns in retinal and cerebral development: radial and tangential.^50^ Tangential migration means that neurons follow the axis perpendicular to the tissue’s apico-basal axis (XY plane), but this rarely occurs in the vertebrate retina,^51^ where most neurons and progenitors travel radially. RGCs are the first-born retinal neurons in the retina, and after their separation from the intermediate committed precursor, they migrate to the most basal layer adjacent to the lens (Z axis). Confocal tracing of individual RGCs shows that this happens through somal translocation (ST) during development.^52^ ST involves a neuron sending their process towards their final location and translocating their cell body along that process.^53^ A challenge in extrapolating these findings to cell transplantation in a developed eye is the lack of developmentally relevant chemokine gradient in the mature retina. The regeneration and renewal of RGCs in lower vertebrates may be more relevant for the transplantation setting. In zebrafish, RGCs can move through multipolar (MP) migration with several processes extending from a cell;^54^ however, it is not known if such a pattern exists in mammalian RGCs.

We performed cluster identification using published single-cell RNA sequencing data to define each retinal neuron population (Fig. 2A-D). RGCs were then subclustered based on their expression of mature (RBMPS) and immature (Pou4f2) markers (Fig. 2A-D, feature plots). We quantified the gene expression changes associated with MP and ST migration and performed a gene set enrichment analysis (escape package)^41,42^ on RGCs from these different developmental stages. During early development, both expressions of the genes related to neuronal migration were more highly expressed, but as RGCs matured (FD82, RBMPS+), expression levels for these genes decreased (Fig. 2E) – indicating that RGC migration occurs early in development and primarily for immature RGCs. Further analysis revealed that genes corresponding to MP migration and ST have different expression patterns, with MP migration genes having their highest expression later in development than ST genes (Fig. 2F). These results are consistent with the fact that MP migration is the preferred migratory mode for late-born neurons that need to navigate through more complex tissue and cellular structures.^24,25^ Therefore, the MP migration and ST gene expression patterns shown here are the first indirect evidence of different modes of RGC migration within the human retina during development. However, despite the evidence of host RGC migration, these data do not indicate how neuron migration is controlled in the retina.

**Fig. 2.**
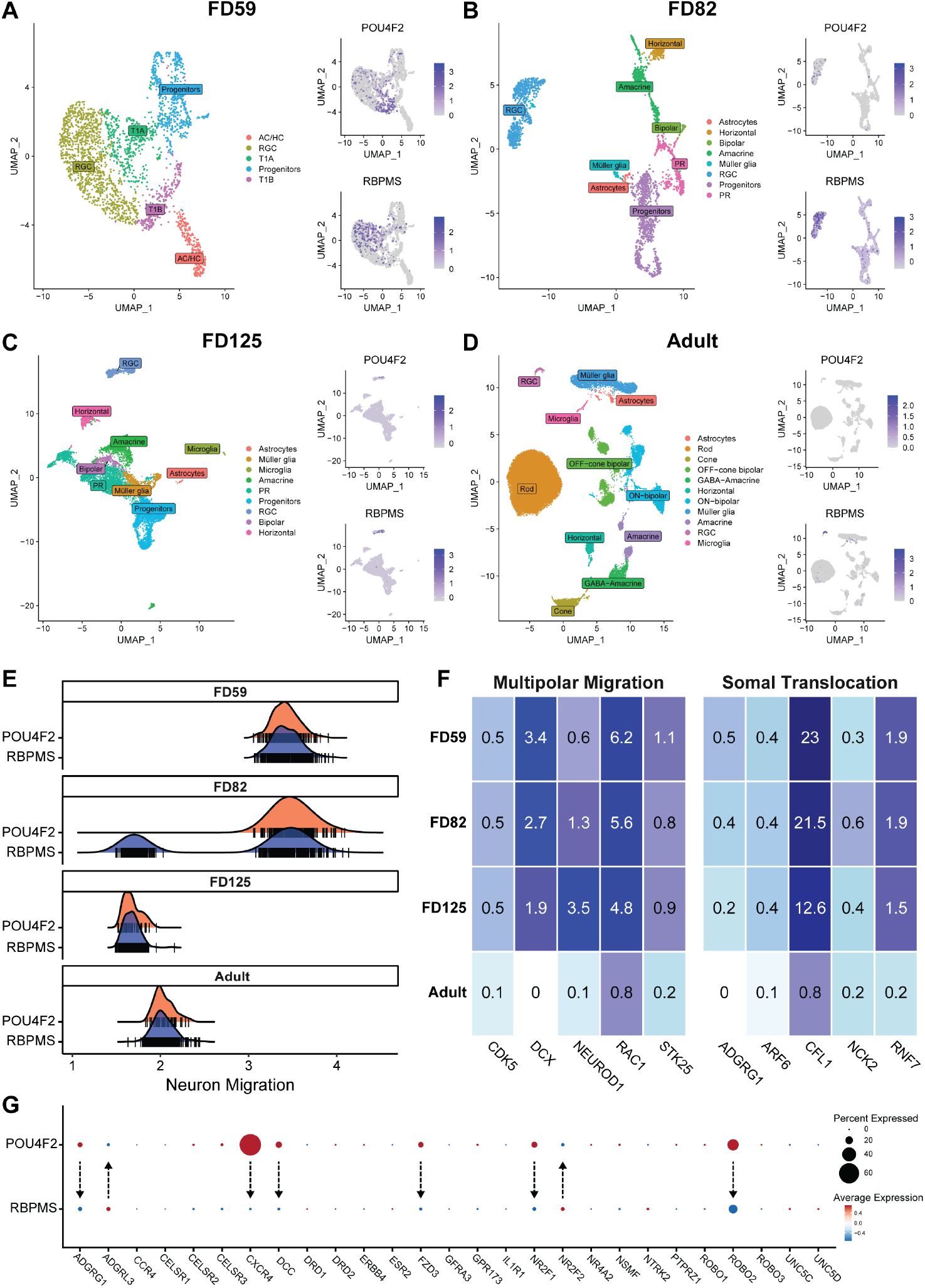
Characterization of human RGC migration patterns and receptor expression during development. **(A)** UMAP plot of the human neural retina with clusters identified by cell-type-specific gene expression and feature plots showing characteristic genes describing clusters of mature (RBPMS) and immature (Pou4f2) RGCs during fetal day 59, **(B)** fetal day 82, **(C)** fetal day 125, and **(D)** adulthood. **(E)** Escape analysis of mature and immature RGC populations during different developmental ages shows the upregulation of genes associated with migration early in development and in immature RGCs. **(F)** Heat map comparison of the most highly expressed genes related to multipolar migration and somal translocation shows somal translocation has the highest expression of these genes during early development (FD59), while multipolar migration has the highest expression of these genes in later stages of development (FD82-125) with little to no expression during adulthood. **(G)** Dot plot showing the percent and average expression in mature (RBPMS) and immature (Pou4f2) RGCs of all the receptors known to be involved in neural migration. Each arrow shows a decrease in expression for all the receptors expressed in > 5% of RGCs and helps to define which receptors may be critical for early RGC migration during development.

The neuron migration can be mediated through transcription factors, adhesion molecules, ion channels, and extracellular cues/ receptors.^55^ Here, we focused on pro-migratory receptors since these are most amenable for modulation through engineering the microenvironment. We screened all the receptors associated with neuron migration that had higher expression in the Pou4f2+ RGC vs. RBMPS+ RGC subclusters and identified six candidates from this in silico analysis to investigate further: Adhesion G Protein-Coupled Receptor G1 (ADGRG1), CXC chemokine receptor type 4 (CXCR4), Netrin receptor DCC (DCC), Frizzled class receptor 3 (FZD3), Nuclear Receptor Subfamily 2 Group F Member 1 (NR2F1), and Roundabout Guidance Receptor 2 (ROBO2) (Fig. 2G).

### CXCR4 activation by stromal cell-derived factor-1 (SDF1) enhances RGC recruitment in vitro

The human stem cell-derived RGCs were used to investigate our ability to control neuron migration. Human RGCs were differentiated from Brn3b-tdTomato hESC in organoid cultures using a 3D-2D-3D technique to parallel development. By recapitulating development in 3D, we expect RGCs differentiated within retinal organoids to have a receptor expression and migratory potential similar to human RGCs.^33^

During the first 3D phase, embryoid bodies are formed by forced aggregation, anterior neural development promoted by WNT inhibition, and retinal differentiation amplified by BMP4 signaling (Fig. 3A). Optic vesicles from the resulting organoid were dissected by random chopping and allowed to attach to a 2D surface to generate an epithelial cell layer (Fig. 3B). During this epithelial formation, retinal progenitor expansion was enhanced by sonic hedgehog activation. Retinal aggregates were then transitioned back to 3D culture and maintained in long-term retinal differentiation media (LT-RDM) that was later supplemented with all all-trans retinoic acid to increase the survival and differentiation of RGCs (Fig. 3A). RGC were isolated from organoid cultures using enzymatic dissociation and magnetic bead sorting against CD90.2 after 42 – 50 days of differentiation. For every stem cell used to form embryoid bodies initially, our protocols resulted in an average of one RGC. RGCs differentiated from Brn3b-tdTomato hESC were visualized by the expression of the tdTomato reporter (Fig. 3B).

**Fig. 3.**
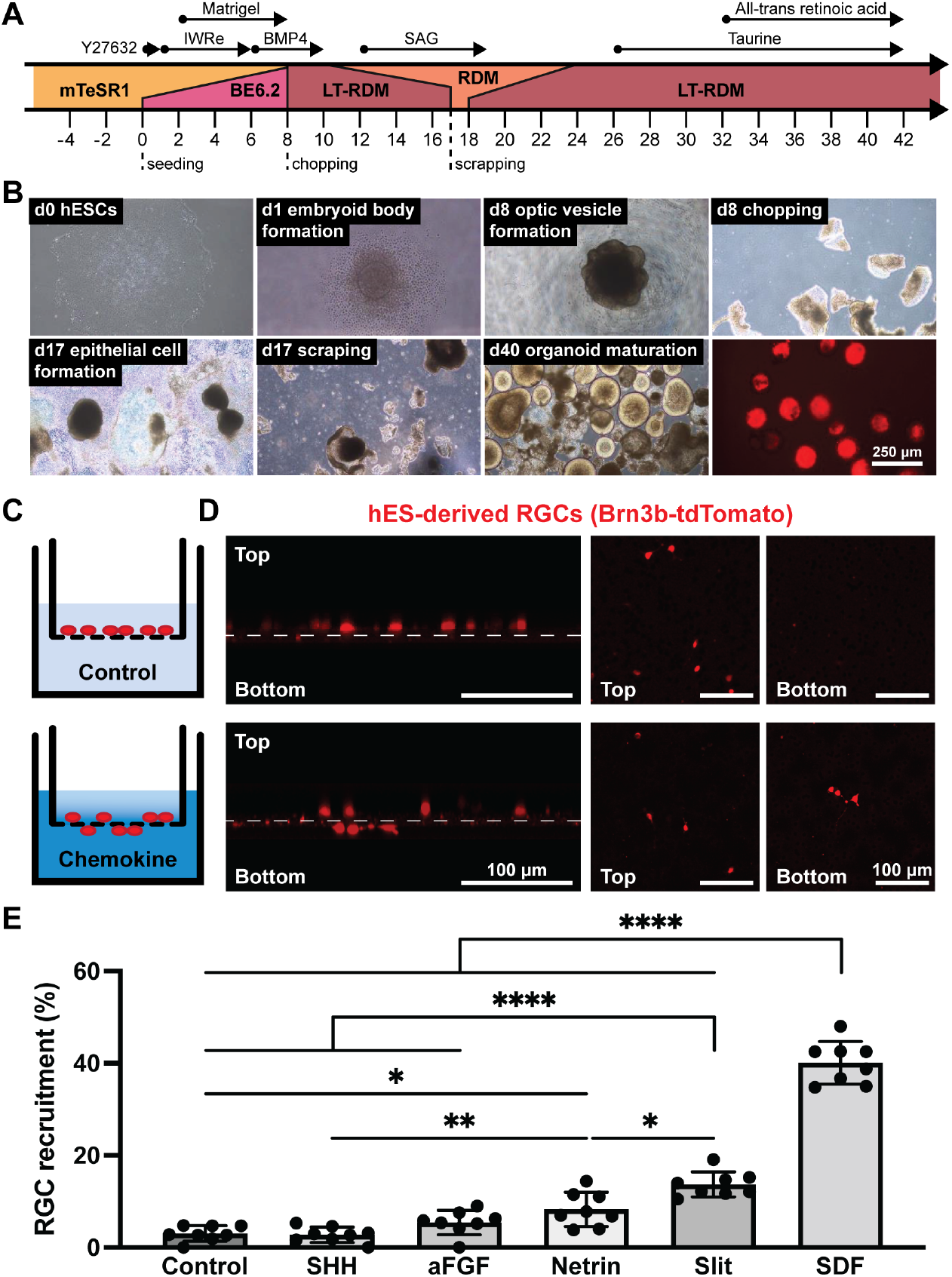
Human stem cell-derived RGCs migrate in response to chemokines. **(A)** Schematic overview summarizing hESC differentiation into retinal organoids using a 3D-2D-3D technique adopted from the Meyer and Zack labs.^26,27^ **(B)** Representative images of important stages in RGC differentiation starting from hESCs to form embryoid bodies and optic vesicle-like structures in 3D culture before being moved to 2D culture to enable epithelial cell formation and then back to 3D culture to mature and generate RGCs (tdTomato positive cells). **(C)** Schematic of cell recruitment transwell assay and **(D)** representative immunofluorescent 3D reconstruction and en-face images of hRGCs on the apical and/or basal membrane surface following chemokine treatment. **(E)** Quantitative analysis of RGC recruitment, defined as the ratio of RGCs on the basal surface to the total number of RGCs, in response to 500 ng/mL SHH, 250 ng/mL aFGF, 100 ng/mL Netrin, 1000 ng/mL Slit, and 50 ng/mL SDF1 treatment. N = 8 wells per group.

Based on our single-cell RNA sequencing analysis and previous cell migration studies,^56^ we selected a panel of chemokine receptors that could enhance and even control neuron cell migration. From our identified receptors, ADGRG1 was excluded because it is a negative regulator of cell migration,^57^ FZD3 was excluded because its ligand, WNT is a key regulator of RGC differentiation,^27^ and NR2F1 was excluded because its ligand is currently unknown.^58^ Fibroblast growth factor receptor (FGFR) was included as a positive control based on prior studies,^59,60^ and Protein patched homolog 1 (PTCH1) included as a negative control because sonic hedgehog protein (SHH) is known not to affect migration. We confirmed the expression of our panel of chemokine receptors in the developing retina and observed different expression levels in each neuron population at each time point (Fig. S1). Critically, in addition to endogenous expression in the neural retina, we also confirm the expression of these receptors in our stem cell-derived RGCs: acidic fibroblast growth factor (aFGF), Netrin1, SHH, Slit1, and SDF1 (Fig. S2A).

To identify our lead candidate for RGC recruitment, we studied the migration of human RGC in response to different chemokine gradients in vitro. We evaluated the effect of aFGF, Netrin1, SHH, Slit1, and SDF1 on RGC recruitment using a transwell assay (Fig. 3C). Chemokine concentrations were selected based on prior work, and a non-treated control was included as a negative control. RGCs were seeded on the apical surface of the membrane and migrated through the pores (8 microns) over 24 hr. RGC were counted in the same areas on both sides of the membrane, and RGC recruitment was calculated as the ratio of cells on the basal surface to the total number of cells (Fig. 3D). Netrin1, Slit1, and SDF1 treatment significantly increased cell recruitment compared with the control, whereas aFGF and SHH showed no statistically significant differences compared to the non-treated control (Fig. 3E). SDF1 led to the most significant increase in RGC recruitment, with 40.11 ± 4.62% of RGCs migrating onto the basal membrane surface. Interestingly, despite being included as a positive control, aFGF failed to cause a significant increase in RGC recruitment, highlighting the power of our approach towards identifying targets to control neuron behavior.

SDF1 and its receptor, CXCR4, are involved in various physiological and pathological processes, development, regeneration, and repair of the nervous system.^61,62^ The role of SDF1/CXCR4 in the retina has been extensively studied in vascular development^63,64^ and pathology.^65–69^ SDF1/CXCR4 is essential in retinal lamination, RGC and photoreceptor development,^70^ and axon regeneration.^61^ Exogenous SDF1 has recently been considered for therapeutic application with profound neuroprotective effects on photoreceptors in animal models of retinal detachment, supported by clinical sample studies with a confirmed elevation of SDF1 in human retinas after detachment.^71^ The vascular effects have to be considered in future clinical applications, and although the role of SDF1 in the progression of retinopathy is described,^72^ there is no evidence to support its role in choroidal neovascularization or Age-related macular degeneration initiation.^73^ The pro-survival effect of SDF1 has been extensively described^74^ with the effect mediated through MAP kinases ERK1/2 and p38. SDF1 and CXCR4 are also required for RGC survival and axon guidance in mammalian and zebrafish retinal development.^75–77^ CXCR4 overexpression in retinal pigment epithelium leads to migration in response to SDF1.^78^ These roles for SDF1 have been explored for axon and optic nerve regeneration,^79^ but not yet explored in the transplantation setting to stimulate RGC recruitment, although CXCR4 is expressed on hESC-derived RGCs.^80^

After identifying SDF1 as the most potent chemokine for RGC recruitment, we sought to better characterize and understand this mechanism. To further validate CXCR4 expression in our RGCs, we performed an immunohistochemistry analysis on culture RGCs. While CXCR4 seems to be expressed in all tdTomato positive cells (Fig. S2B), it is also expressed in tdTomato negative cell populations (Fig. S2C, yellow arrow). We confirmed this result by flow cytometry and single-cell RNA sequencing of retinal organoids. Flow cytometry shows that greater than 85% of all tdTomato positive cells (hRGCs) express CXCR4 by flow cytometry (Fig. S2D). Similarly, by single-cell RNA sequencing, we show that greater than 55% of the RGC cluster express CXCR4 (Fig. S3). Interestingly, while most hRGCs express CXCR4, only ∼40% migrated in response to SDF1 treatment. Therefore, we again adopted our transwell assay to evaluate the effect of SDF1 concentration on RGC recruitment to understand if this response is dose-mediated. Using the same approach, we evaluated various concentrations of SDF1: 0, 10, 50, 200, and 500 ng/mL. When compared to the control, 10, 50, and 200 ng/mL SDF1 resulted in significantly enhanced RGC recruitment, but there was no statistically significant difference between 50 and 200 ng/mL SDF1 (Fig. S4A, 39.71 ± 5.67 vs. 45.85 ± 7.52), indicating some upper limit to this dose-response. No RGCs remained attached to the membranes in cultures treated with 500 ng/mL SDF (not shown). Lastly, we investigated the ability to inhibit SDF1-mediated RGC recruitment by the preconditioning donor RGCs with a CXCR4 antagonist, AMD3100. Blocking CXCR4 hRGCs with 10 µM AMD3100 in our transwell assay resulted in complete inhibition of RGC recruitment by SDF1 treatment (Fig. S4B, 42.18 ± 10.48 vs. 12.73 ± 1.81).

### An SDF1 gradient across the neural retina improves the structural integration of donor RGC in vivo

After demonstrating that SDF1 treatment enhances RGC recruitment in vitro (Fig. 4E, Fig. S4A), we transplanted human RGCs subretinally and delivered recombinant SDF1 protein intravitreally to establish a chemokine gradient across the retina and drive donor RGC migration in vivo (Fig. 4A). To deliver donor RGCs, we accessed the SR space between the retinal pigment epithelium and photoreceptor outer segments through the sclera to avoid causing a retinotomy that could limit the effectiveness of our chemokine gradient. Furthermore, to support donor RGCs within the SR space and establish a proneuronal microenvironment like that of our cell culture system, we formulated our donor cells with slow-release neurotropic factors (GDNF-, BDNF-, and CNTF-loaded polyhedrin-based particles (PODs)). Lastly, unlike our mouse syngeneic transplantations (Fig. 2A), retinas were collected for analysis three days following cell delivery to limit the immune response in our xenotransplantation experiments.

**Fig. 4.**
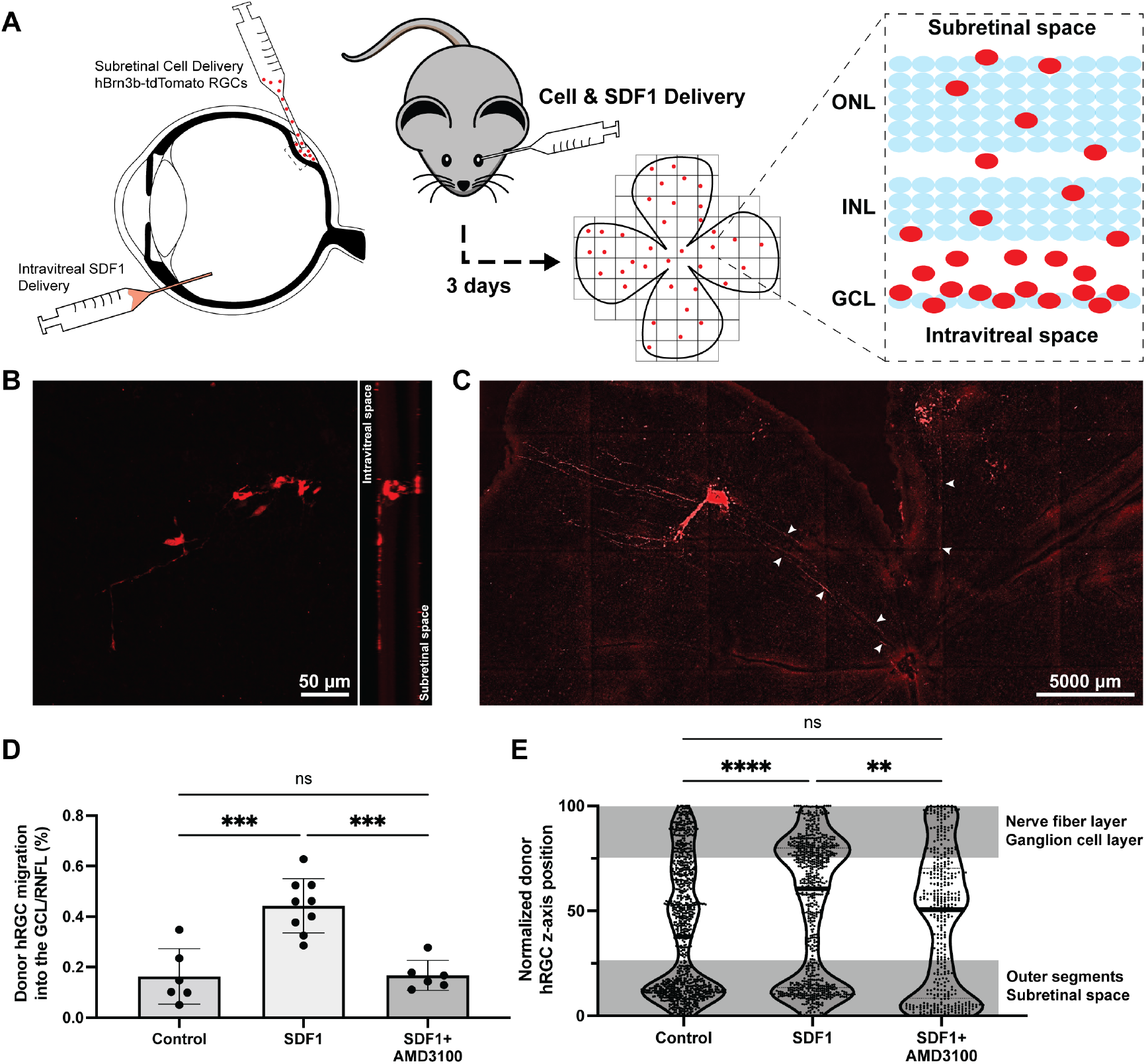
Human stem cell-derived RGC xenotransplantation in mice with exogenous SDF1 gradient. **(A)** Schematic overview of our donor RGC and SDF1 delivery strategy to establish a chemokine gradient across the neural retina and quantification paradigm to assess structural integration of individual donor RGCs 3 days following transplantation using a 3D reconstruction of a retinal flat mount. **(B)** Representative en-face immunofluorescent image (max intensity projection) and orthographic projection showing donor RGC (red) migration from the subretinal space towards the intravitreal space in response to SDF1 treatment. **(C)** Representative retinal flat mount (max intensity projection) shows donor RGCs extending neurites towards the optic nerve head (white arrows) following SR cell delivery and IVT SDF1 treatment. **(D)** Quantitative analysis of donor RGC migration into the GCL/RNFL shows that the significant increase in donor RGCs translocating into the GCL/RNFL in response to SDF1 treatment is mediated through CXCR4. N = 6 – 8 mice per group. **(E)** Single-cell quantification of z-axis position normalized to the thickness of the retina reveals that while SDF1 treatment enhances the structural integration of donor RGCs, a significant population of donor RGCs fail to migrate and remain within the subretinal space. The nerve fiber layer and ganglion cell layer were defined as the top quartile of the retina, while the subretinal space and outer segments were defined as the bottom quartile.

Retinal explants were prepared as flat mounts and imaged using confocal microscopy to assess donor RGC distribution (Fig. 4B). The resulting tiled, z-stacked image was reconstructed using custom semi-automated ImageJ and MATLAB scripts to identify the x-, y-, and z-position of each donor RGC within the neural retina. The z-axis position was then normalized to the thickness of each retina so those values could be superimposed and differences between experimental groups evaluated. Due to differences in each retinal preparation and the difficulties defining the retinal nerve fiber layer (RNFL) from the GCL and the outer segments from the SR space, we defined the GCL and RNFL and the SR space and out segments together as the top and bottom quartile of the retina, respectively.

Previously, BDNF was shown to enhance the migration of sensory neurons in zebrafish,^81^ but its role in neuron migration in the transplantation setting remains unknown. Therefore, before studying how SDF1 affects RGCs, we first investigated the role of PODs on migration in vivo. While the PODs formulation resulted in increased neurite outgrowth (Fig. S5A), it did not have any apparent effect on donor RGC translocation into or towards the GCL (Fig. S5B-C).

Following SR delivery of donor RGCs, we observed both spontaneous and chemokine-directed migration within the neural retina into the GCL/RNFL (Fig. 4B). Critically, because the ILM was left intact during our transplantations, this membrane could function as a stop signal, and we did not detect any cells on the vitreous surface of the retina. Moreover, we observed neurite outgrowth towards the optic nerve head in a subset of transplantations from those that received SDF1 (Fig. 4C, white arrows). SDF1 treatment also significantly increased RGC migration into the GCL/RNFL from 16.4

± 11.0% to 44.3 ± 10.8% (Fig. 4D) – a 2.7-fold increase in migration into the GCL/RNFL. Approximately the same percentage of donor RGCs (∼ 40%) migrated in response to SDF1 treatment in both an in vitro and in vivo environment. We do not yet know if other intrinsic cell characteristics outside CXCR4 expression result in this similar response or if additional SDF1 injections will further enhance RGC recruitment.

To better understand this response and to demonstrate a direct effect of SDF1 on donor RGCs rather than some secondary effect, we preconditioned donor RGCs with AMD3100 to block the CXCR4 receptor. Blocking SDF1 binding to the CXCR4 receptor with AMD3100 inhibited SDF1-mediated donor RGC translocation into the GCL/RNFL (Fig. 4D). Despite preventing migration into the GCL/ RNFL, blocking CXCR4 activation with AMD3100 does not appear to completely inhibit donor RGC migration in response to SDF1, with a discrete population of donor cells in the central layers of the retina (Fig. 4E). Furthermore, while SDF1 treatments significantly increased the number of donor RGCs within the GCL/RNFL (Fig. 4D), a large population of donor RGCs failed to migrate in response to SDF1 and remain within the SR space (Fig. 4E). No significant differences in total donor RGCs were detected between groups (Fig. S6A).

In a final set of transplantation experiments, we aimed to validate chemokine-directed donor cell migration across species and in different directions. To demonstrate this phenomenon, we delivered mouse stem cell-derived RGCs SR or IVT and injected SDF1 IVT or SR to establish a forwards and reverse gradient across the retina, respectively (Fig. S7A-C). Two weeks after transplantation, retinas were stained for host and donor RGCs to assess donor cell integration and distribution across and within the retina with respect to the host GCL. Given the longer possible timeframes for syngeneic transplantation, donor RGC morphology was also analyzed to assess their capacity to extend their processes towards the inner plexiform layer (IPL). For both SR and IVT cell delivery, the artificial SDF1 gradient increased donor RGC migration/neurite extension in the direction of SDF1 and the amount of donor cell processes found within the IPL (Fig. S7B-D). These results further highlight that SR cell delivery improved RGC integration into the host retina compared to IVT cell delivery, irrespective of SDF1 treatment. To provide one final proof of concept study, we transplanted intact retinal organoids (Fig. S8A) to the SR space and delivered SDF1 IVT. Quantifying normalized donor cell distribution shows donor RGC migration/neurite extension out from the retinal organoids in the direction of our SDF1 treatment (Fig. S8B).

### Human retinal neurons primarily migrate through the multipolar (MP) migration mode following transplantation in mice

To understand the mechanisms by which donor RGCs migrate through the mature retinal tissue in response to SDF1 treatment, we first investigated RGC migration patterns and kinetics in vitro. We confirmed the capacity for stem cell-derived RGCs to migrate via each migratory modality within 3D tissue by live cell imaging in mouse retinal organoids. Neurons migrating via ST will extend a single neurite towards their final location and then pull their soma along that neurite with or without retracting it (Fig. S9A). Conversely, neurons migrating via MP migration will extend multiple processes in different directions as they pull their soma towards their final location in a dynamic but random pattern (Fig. S9A). Following translocation, RGCs lose their remaining processes and project axons toward the optic nerve. To visualize neuronal migration within retinal organoids, RGCs were differentiated using a sparse Thy1.2-GFP stem cell line (Fig. S9B). Critically, RGCs differentiated from this reporter line express green fluorescent protein (GFP) fluorescence within their processes, which can be observed to confirm each migratory mode. Qualitative observation of individual frames in series demonstrates that mouse stem cell-derived RGCs can migrate by ST (Fig. S9B’) and MP migration (Fig. S9B’’). While ST was the most common migratory mode observed within the retinal organoids, these confocal traces of individual neurons represent the first demonstration of spontaneous MP migration for mammalian stem cell-derived RGCs.

However, we could not rely on spontaneous migration to confirm these migratory modes in vivo because of our limited capacity to quantify live cell migration in vivo. Moreover, unlike mouse retinal organoids, due to the dense and robust fluorescent reporter expression within retinal organoids derived from Brn3b-tdTomato hESCs, we could not quantify migration within 3D tissues in vitro. Therefore, we utilized small molecule migration inhibitors to investigate the migration modes of human RGCs in vitro and in vivo. To confirm the efficacy of these inhibitor molecules and study each migration pattern, RGCs were differentiated in 3D organoid cultures, isolated by magnetic microbeads against CD90.2, and cultured on laminin-coated plates for live cell video microscopy. We temporarily destabilized the cytoskeleton of donor RGCs to alter their migratory modality between ST and MP migration or completely inhibit cell motility. By treating RGCs with a CDK5 inhibitor (roscovitine, 15 µM),^82,83^ MP migration is inhibited, and ST will be the only active migratory mode (Fig. S10A). Similarly, by treating RGCs with an Arp2/3 inhibitor (CK-666, 200 µM),^84,85^ ST is inhibited, and RGCs will only be capable of migration via MP migration (Fig. S10B). RGC viability was > 90% for each group, indicating that each inhibitor molecule did not affect cell viability (Fig. S10C). Each migration pattern was confirmed with an in vitro time-lapse study with 20 min intervals. Inhibiting ST with CK666 resulted in a significant decrease in RGC migration speed and total displacement from 5.6 ± 2.2 to 4.7 ± 2.0 µm/s and 160.1 ± 85.0 µm to 139.8 ± 71.6 µm, whereas inhibiting MP migration with roscovitine decreased their speed to 3.2 ± 1.5 µm/s and 93.9 ± 54.0 µm, respectively – demonstrating hESC-derived RGCs can migrate via both modalities (Fig. S10D-F). To further validate these results, we showed that the coadministration of both inhibitor molecules prevented migration in most treated cells in vitro (Fig. S10D-F). Somewhat surprising is that ST inhibition by roscovitine treatment resulted in significantly slower RGC migration than MP migration inhibition by CK666 treatment, despite ST being the faster mode of migration. Furthermore, the average speed and displacement of cells traveling by each of the two migration modes do not sum to the average migration kinetics of the non-treated controls. This finding indicates that either there are some off-target effects of roscovitine that further limit migration or not all RGCs have the same capacity to migrate by each mode. The discrete cell populations observed in the violin plots suggest the latter explanation is more likely (Fig. S10D-E) while also demonstrating the effectiveness of each small molecule in inhibiting each mode of migration independently.

By applying these same inhibitor molecules to our in vivo transplantation paradigm, we demonstrated distinct patterns of donor RGC coverage and positioning within the neural retina (Fig. 5A-D). Our results show impairing ST by CK666 preconditioning does not affect the percentage of donor RGCs that migrate into the GCL/RNFL in response to SDF1 (No inhibition: 44.3 ± 10.8%; ST inhibition: 47.2 ± 12.0%, Fig. 5E). However, inhibition of MP migration by roscovitine preconditioning significantly limits the percentage of donor RGCs that migrate into the GCL/RNFL in response to SDF1 compared to the CK666 preconditioning and no treatment groups (MP migration inhibition: 20 ± 3.7%, Fig. 5E). Critically, each mode of neural migration inhibition did not affect the total number of surviving donor RGCs (Fig. S9A). The fact that there were no significant differences between dual inhibition of migratory modes and MP migration inhibition alone (dual inhibition: 14.7 ± 8.1%, Fig. 5E) indicates that MP migration is the primary mode by which donor RGCs migrate within the neural retina in response to SDF1 treatment. Despite no significant difference in the number of donor RGC structurally integrating into the GCL/RNFL following SDF1 treatment between the control donor RGCs and those preconditioned with CK666, CK666 preconditioning resulted in a significant increase in overall migration towards the GCL/RNFL (Fig. 5F). However, while CK666 preconditioning increased the overall migration towards the GCL/RNFL, we do not yet know why these donor RGCs failed to continue into the GCL/RNFL.

**Fig. 5.**
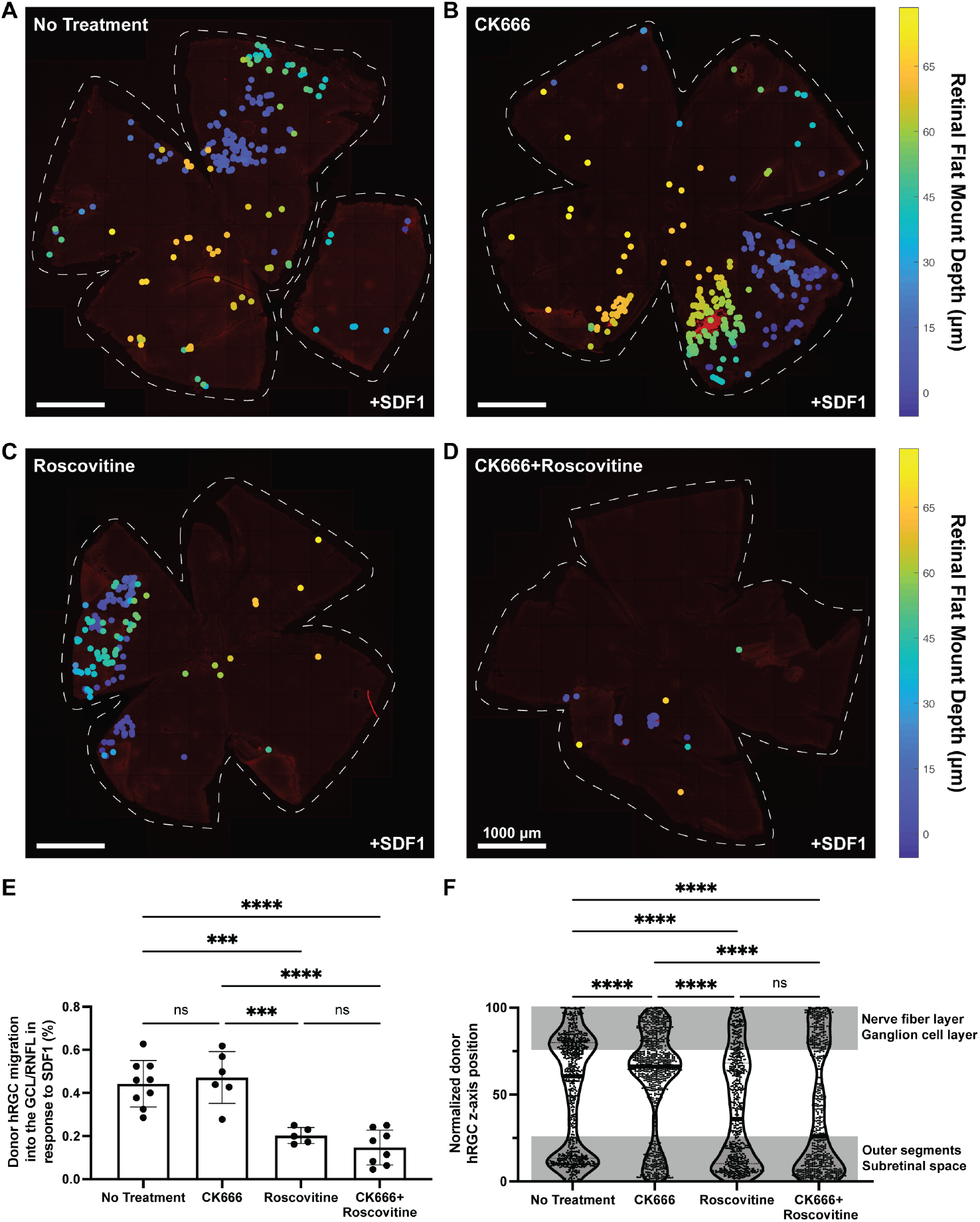
Distinct patterns of donor neuron translocation following SDF1 treatment in mice. **(A)** Representative depth-coded color map overlayed on retinas transplanted with donor RGCs preconditioned with DMSO (no treatment control), **(B)** CK666, an ST inhibitor, **(C)** roscovitine, an MP migration inhibitor, and **(D)** CK666 + roscovitine to modulate their migratory modality shows donor RGC distribution and coverage across and within the retina following SDF1 treatment. Deep blue represents the subretinal space, while yellow represents the GCL/RNFL. **(E)** Quantitative analysis of donor RGC migration into the GCL/ RNFL in response to SDF1 shows no difference between control donor RGC and those preconditioned with CK666, but a significant decrease in translocation for donor RGC treated with roscovitine and CK666 + roscovitine, indicating MP migration, but not ST, is required for donor RGCs to migrate into the GCL/RNFL through the neural retina. N = 5 – 8 mice per group. **(F)** Single-cell quantification of z-axis position normalized to the thickness of the retina reveals that while there was no significant difference in the number of donor RGC structurally integrating into the GCL/RNFL between the control donor RGCs and those preconditioned with CK666, CK666 preconditioning resulted in a significant increase in overall migration towards the GCL/RNFL. The nerve fiber layer and ganglion cell layer were defined as the top quartile of the retina, while the subretinal space and outer segments were defined as the bottom quartile.

RGCs are also capable of tangential migration due to retinal refinement during development and age-related degeneration,^86,87^ yet we see limited migration in the x- and y-axes following transplantation. Unfortunately, SDF1 treatments alone were insufficient to significantly increase donor RGC retinal coverage, defined as the percentage of tiles per retina with one or more donor RGC (Fig. 5B). However, in combination with ST inhibition by CK666 preconditioning, donor RGC retinal coverage was significantly increased from 23.6 ± 7.8% for donor RGCs without preconditioning, and no SDF1 injections to 50.5 ± 27.5% for donor RGCs preconditioned with CK666 and with SDF1 injections (Fig. 5B).

### SDF1 treatment causes the displacement of endogenously regenerated retinal neurons in situ

Despite our CXCR4 blocking experiment demonstrating the direct effect of SDF1 on donor RGC translocation into the GCL, host neurons within the retina also express CXCR4,^88^ and could be adversely affected by SDF1. To investigate the effects of SDF1 on retinal lamination and position of host retinal neurons, we delivered a range of concentrations (0, 1, 10, and 200 ng SDF) into the mouse vitreous and collected retinas for histology 24 hours after the injection. Hematoxylin and eosin (H&E) stained sections of the neural retina following intravitreal injection SDF1 showed no apparent changes in retinal lamination (Fig. S11A). Retinal sections were also stained for host neurons and glia (RGCs, bipolar cells, photoreceptors, and Müller glia) to visualize any effects of SDF1 treatments. We observed no differences in RGC, bipolar cell, and photoreceptor morphology and position between sections and concentrations of SDF1 (Fig. S11B-D), but Müller glia (MG) appeared to be reacting to SDF1 treatments (Fig. S11E). To quantify this reaction, we calculated the normalized displacement error (NDE), defined as the difference between the total segment length connecting each point and a straight line normalized to the total number of points (Fig. S12). For example, a high NDE will be associated with increased total displacement from the mean. By applying this analysis, we demonstrated that MG have increased displacement with increasing concentrations of SDF1 (Fig. S11F).

While stem cell replacement therapies represent one approach to restoring retinal neurons lost in glaucoma and other optic neuropathies, stimulating neurogenesis in the retina through endogenous reprogramming of MG may be an alternative regenerative strategy. Todd et al. demonstrated efficient stimulation of retinal regeneration from MG in adult mice using a combination of proneuronal transcription factors; however, following neurogenesis, RGC-like neurons mostly fail to migrate to the GCL.^11^ Here, we sought to improve the migration of MG-derived newborn neurons using SDF1 to demonstrate the universality of chemokine-directed migration for regenerative medicine.

MG were reprogrammed into RGC-like neurons as previously described.^11^ Approximately one week after initiating reprogramming, host RGCs were ablated to mimic the loss of RGCs in glaucoma using NMDA-inducted toxicity. After a short recovery period (1 – 2 days), SDF1 was injected intravitreally to test whether endogenously regenerated neurons can be recruited by SDF1. Two weeks following SDF1 treatment, the locations of MG-derived neurons (GFP+/HuC/ D+) were quantified (Fig. 6A-B). We found that SDF1 elicited a dose-dependent displacement of MG-derived neurons. Newly generated neurons showed an NDE from 3.59 ± 1.50 to 11.8 ± 5.33 at 0 and 100 ng SDF1 delivery, respectively (Fig. 6C) – a 3.3-fold increase in displacement. These results highlight the potential to direct reprogramed neuron migration in situ with chemokines. In addition to regenerative medicine, repositioning MG-derived neurons in vivo may also enable improved access for quantitative assessment, such as whole cell patch-clamp recordings. However, we do not yet know the ideal timing for SDF1 treatment nor if displacement occurs before or after reprogramming. The delivery of a slow-release formation of SDF1 may help to solve this problem in the future and lead to enhanced neural recruitment for both endogenously reprogrammed and stem cell-derived donor neurons.

**Fig. 6.**
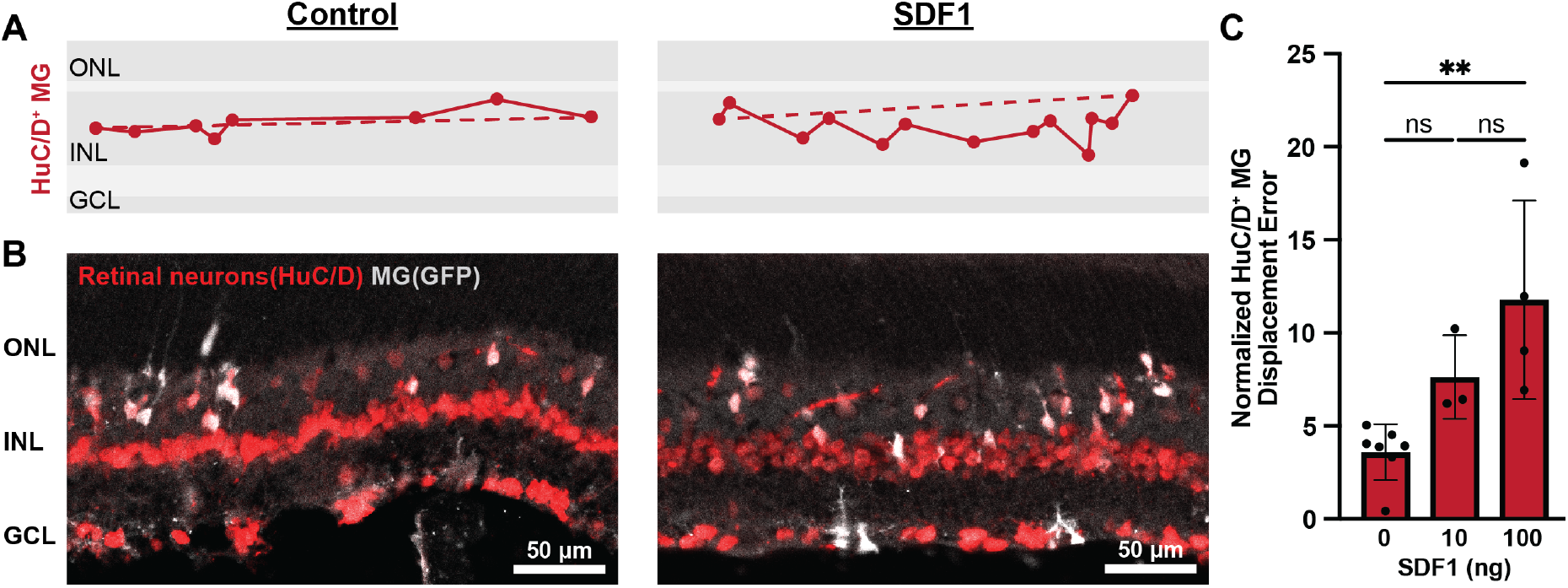
Endogenous reprogramed neurons response to SDF1 treatment in the retina. **(A)** Representative quantitative assessment of normalized displacement error showing the positions of HuC/D positive MGs (red and white colocalization) and **(B)** immunofluorescent images of retinal sections stained for MGs (GFP positive, pseudo colored gray) and retinal neurons (HuC/D positive, red) with and without SDF1 treatments. **(C)** Quantification of normalized displacement error for HuC/D positive MGs in an NMDA damaged model at 0, 10, and 100 ng SDF1 shows a dose-dependent response. ONL: outer nuclear layer; INL: inner nuclear layer; GCL: ganglion cell layer.

## Conclusions

The transplantation of stem cell-derived dopaminergic neurons, motor neurons, photoreceptors, retinal pigment epithelium, mesenchymal stem cells, and other cell types has reached the stage of clinical trials. The preclinical studies of cell engraftment into various animal models highlighted the importance of the niche and the need to better control cell behavior after delivery by engineering cell-intrinsic and extrinsic factors. Here, we describe the framework to identify microenvironment cues to control a specific donor cell behavior using retinal ganglion cells (RGCs) in the eye as a substrate and migration as a function of interest. This “in silico – in vitro – in vivo” funnel can be applied to other cellular processes, including synapse formation, phagocytosis, insulin production, axon growth, etc.

Moreover, identifying SDF1 as a chemokine, leading to a > 2-fold increase in donor cell migration, allowed us to study the migration of neurons in the controlled and accessible ecosystem of the eye. We have demonstrated that RGCs can migrate via somal translocation and multipolar (MP) migration modes in vitro; however, they primarily use MP migration in vivo in the adult retina to navigate the developed tissue architecture. Donor RGC migration allows us to direct donor cells within the ganglion cell layer within the first few days after transplantation, allowing them to integrate into existing retinal circuitry.

## Methods

### Mouse cell culture and mouse RGC differentiation

Mouse iPSCs derived from Tg[Thy1.2-Green Fluorescent Protein (GFP)]M mouse fibroblasts were maintained in gelatin-coated (0.2% v/v) tissue culture flasks in standard culture conditions (5% CO2, >95% humidity, 37°C). Mouse stem cell maintenance media [10% fetal bovine serum (FBS), 1x L-glutamine, 1x non-essential amino acids, 1x sodium pyruvate, 1x antibiotic-antimycotic, 1x EmbryoMax Nucleosides, 100 μM b-Mercaptoethanol, and 100 U/mL mouse leukemia inhibitory factor in DMEM/F12 (3500 mg/L dextrose)] was exchanged every other day until colonies reached ∼70% confluency. After reaching ∼70% confluency, iPSCs were collected with 1x trypsin-EDTA as a single-cell suspension to seeded new gelatin-coated tissue culture flasks at 2000 cells/cm^2^ or to differentiate RGCs as previously described.^29^ In brief, spheroids were formed by seeding 1500 cells in 50 μL OV media [Optic vesicle media: 1x L-glutamine, 1x non-essential amino acids, 1x sodium pyruvate, 1x antibiotic-antimycotic, 1x lipid concentrate, 0.2x Insulin-Transferrin-Selenium-Ethanolamine, 100 μM b-Mercaptoethanol, 1.5% FBS, 1 μM N-Acetyl-L-cysteine (NAC), and 10 μM forskolin in DMEM/F12 (3500 mg/L dextrose)] per well in uncoated polystyrene U-bottom 96-well plates. After 24 hours at standard culture conditions, an additional 50 μL OV media supplemented with 2% Matrigel was added to each well to induce forebrain/retinal differentiation. The following day, an additional 100 μL OV media with 1% Matrigel was added to each well, and retinal organoids were maintained in this media until day 9 of differentiation by replacing half the media every other day. On day 9, organoids were removed from the well plate and randomly chopped to dissect the optic vesicles to allow for greater expansion. The resulting retinal aggregates were transferred to a low-attachment culture flask, and were maintained in OC media [Optic cup media: 1x L-glutamine, 1x non-essential amino acids, 1x sodium pyruvate, 1x antibiotic-antimycotic, 1x lipid concentrate, 100 μM b-Mercaptoethanol, 1x B27 without vitamin A, 1 μM N-Acetyl-L-cysteine (NAC), and 10 μM forskolin in DMEM/F12 (3500 mg/L dextrose)] until day 21 – 25 of retinal maturation, with half media changes every other day. From day 16 onwards, the OC media was supplemented with 250nM all-trans-retinoic acid.

### Maintenance of human stem cells and human RGC differentiation

H9-BRN3B:tdTomatoThy1.2-hESCs colonies were maintained in Matrigel-coated (2%) six-well plates with mTeSR1 media in a low oxygen environment (5% CO2, 3.5% O2, >95% humidity, 37°C).30 Media was changed daily with room temperature mTeSR1 media. At 70% confluency, after approximately 4-5 days of culture, hESCs were enzymatically passaged with accutase for 8 min at room temperature and split at a ratio of 1:10 to 1:20 into a new Matrigel-coated six-well plate.

Human RGCs were differentiated from hESCs in 3D retinal organoid cultures using established protocols with minor adaptations.^27,28^ In brief, while passaging hESCs, colonies were enzymatically dissociated for 4 additional minutes (12 minutes total) to yield a single-cell suspension. Embryoid bodies were first formed by aggregating 7,500 hESCs/well with ultra-low attachment U-bottom 96-well plates in a low oxygen environment for 24 hours. The plate was transferred to a standard culture environment after the first 24 hours and the culture media was gradually transitioned from 3:1 mTeSR1:BE6.2-NIM [<u>B</u>27 + <u>E6</u> at <u>2</u>X concentration – neural induction medium: 1x L-glutamine, 1x sodium pyruvate, 1x antibiotic-antimycotic, 1x B27 without Vitamin A, 0.88 mg/mL NaCl,

38.8 mg/L insulin, 128 mg/L L-ascorbic acid, 28 μg/L selenium, 21.4 mg/L transferrin, and 38.8 mg/L NaHCO3 in DMEM/F12 (3500 mg/L dextrose)] to >90% BE6.2-NIM by day 8. Media was exchanged daily until day 6 and was supplemented with 20 μM Y-27632 on day 0, 1% Matrigel on days 2 – 6, 3μM IWR-1-endo on days 1 – 4, and 55 ng/mL BMP4 on day 6 to promote neural development and early retinal differentiation. On day 8 of differentiation, organoids were collected and randomly chopped to excise the optic vesicles to allow for better propagation and maintaining the size below the diffusion limit, predicted to be maximum diameter of 1.4 mm for cerebral organoids.^31^ The chopped aggregates were seeded in LT-RDM [Long-Term Retinal Differentiation Media: 1x L-glutamine, 1x sodium pyruvate, 1x antibiotic-antimycotic, 1x B27, 1x non-essential amino acids, and 10% FBS in 50:50 1x DMEM/F12 (3500 mg/L Dextrose):DMEM (4500 mg/mL dextrose)] on tissue culture treated plastic to encourage attachment and retinal epithelium formation. To enhance retinal progenitor expansion, culture media was exchanged every other day with a gradual transition to 1:1 LT-RDM:RDM [Retinal Differentiation Media: LT-RDM without FBS] by day 17 and supplemented with 100nM smoothened agonist (SAG) on days 12 – 16. On day 17, the retinal tissue aggregates were carefully scrapped and lifted off the tissue culture plastic so their neural epithelium remained attached. The detached retinal aggregates were then transferred to a poly(2-hydroxyethyl methacrylate)-coated flask and cultured in suspension with RDM supplemented with 100nM SAG. The next day, day 18, culture media was replaced with 1:3 LT-RDM:RDM (2.5% FBS) supplemented with 100nM SAG. After day 18, culture media was exchanged every other day and was gradually transitioned to LT-RDM by day 24. From days 26 and 32 onwards, culture media was supplemented with 1mM taurine and 250nM all-trans-retinoic acid, respectively. LT-RDM was exchanged every other day until retinal maturation between days 42 – 50.

### Dissociation and isolation of RGCs

Mature mouse and human retinal organoids were dissociated using the embryoid body dissociation kit (Miltenyi Biotec, 130-096-348) with the gentleMACS™ Dissociator according to the manufacturer’s automated dissociation protocol with a minor modification. Specifically, gentleMACS Program EB_01 and EB_02 were replaced with Brain_01 and Brain_02, respectively. RGCs were isolated from freshly dissociated organoids by magnetic microbead sorting for Thy1.2+ cells using the Dynabeads™ FlowComp™ Mouse Pan T (CD90.2) Kit (Invitrogen, 11465D) according to the manufacturer protocol.

### Single-cell RNA sequencing data analysis and identification of targets

The data analysis was performed using the obtained datasets from Wang et al. for human adult retina,^32^ and Sridhar et al. for fetal (FD59, FD82, FD105) and hPSC-derived retinal organoids (OD45, OD60).^33^

The data remaining after quality control filtering were analyzed in RStudio v. 1.4.1717 (R 4.1.1) using the Seurat package (v. 4.1.0).^34^ The effects of cell cycle heterogeneity in gene expression were mitigated by regressing out the difference between G2M and S phase signatures by Seurat functions CellCycleScoring and ScaleData scaling the gene content by cell (including S/G2M score after cell cycle genes analysis,^35^ RNA count and feature, ribosomal and mitochondrial genes expression). The downloaded human adult retina dataset was pre-integrated, and for FD82 and FD105, peripheral and central retina datasets integration was performed.

The integration steps included rescaling the data, finding the mutual nearest neighbors,^36^ creating the integration anchors, and transforming the datasets during the IntegrateData function with PCs and cluster computations. UMAP was performed,^37^ and the clustering resolution for the datasets were sufficient to match with the cell type markers for the Louvain clustering algorithm with a resolution equal to 0.5.^38^ The markers and their expression level for each cluster as differentially expressed genes (DEGs) were received with FindAllMarkers function (minimum fraction of 0.25 and log2 fold change threshold of 0.25) and further DoHeatmap function for the DEG analysis. The DEGs were also checked for the population labeling consistency in different populations of the Cell Atlas of The Human Fovea and Peripheral Retina.^39^ Clusters with multiple population patterns during the early stages of development were labeled as T (T1A and T1B for FD59) and removed from the analysis. Some clusters with more differentiated but still multiple phenotypes were subsetted and reprocessed with deeper resolution. Then the cells were separated by DEG expression of different known populations using the GetAssayData function and labeled using the SetIdent function.

After the cluster identification was performed and clusters were labeled, the rod and cone, GABA-amacrine and amacrine, OFF-cone bipolar and ON-bipolar were shown separately on the embeddings using DimPlot function but merged to photoreceptors, amacrine and bipolar for the downstream analysis.

Separated expression of different genes was shown using the FeaturePlot function, and all the time points were merged for the gene expression population analysis. The assay objects were created using the CreateAssayObject function by exporting the time point gene expression and generating a new default assay for multiple time points. Then gene expression for each time point was shown using the DotPlot function.

To show the distribution of gene expression, violin plots were generated using the VlnPlot function. RGC population was subsetted for all the time points separately and then merged into a new Seurat object with the following time points: FD59, FD82, FD125, Adult, OD45+60. Organoid datasets for days 45 and 60 were pre-integrated using the same approach. For the percentage of CXCR4+ cells for different time points, a barplot was created using the ggplot2 function.^40^

To check the common migration expression, we used the escape [easy single cell analysis platform for enrichment] package (v. 1.2.0).^41,42^ We integrated the Neuron Migration pathway from Molecular Signatures Database (http://www.gsea-msigdb.org/gsea/msigdb/cards/GOBP_NEURON_MIGRATION.html) for this analysis. Then we merged the RGC datasets separated by POU4F2/ RBPMS for every time point and performed the enrichment on the RNA count data using the enrichIt function with the ssGSEA method.^43^ To examine the enrichment distributions for the neuron migration pathway, we generated a ridge plot using the ridgeEnrichment function.

To assess cell migration pathway gene expression, we separated cell migration into two modes: somal translocation (ST) and multipolar migration (MP). To obtain the gene lists for those groups, we downloaded the Gene Ontology Annotations (http://www.informatics.jax.org/go/term/GO:0001764) and added lists of the well-known genes for those migration pathways appearing in the literature to choose the most significant genes for each mode of migration.^44–47^ The resulting list was given as a vector for features in the DotPlot function. After the genes were chosen, we quantified the average expression of the genes for ST and MP for every time point using the AverageExpression function. The ggplot2 package was used to generate the heatmap with the functions geom_tile, facet_grid applied, and log transformation performed.

To show the differences between immature and mature RGCs, we separated them by POU4F2 (BRN3B) / RBPMS expression and subsetted the resulting populations into new Seurat objects for FD59, FD82, FD125, and adult datasets. The receptor expression for immature and mature RGC populations was shown using the DotPlot function.

The Seurat objects with the annotated UMAP embeddings generated during the study are provided in the Source data file. The packages used for the analysis can be found in Table S1. The code used for this study for reproducing the bioinformatical analysis is available on GitHub at: https://github.com/mcrewcow/BaranovLab

### In vitro cell recruitment assay

The effects of a panel of chemokines (SHH, aFGF, Netrin, Slit, and SDF1) and a CXCR4 antagonist (AMD3100; Calbiochem, 239820) on cell recruitment in vitro were investigated using porous membrane cell culture inserts with 8 μm pore size (Greiner Bio-One PET ThinCerts). The top and bottom of the porous membranes were coated with a 20 μg/mL laminin solution for 4 hours. Excess laminin was rinsed from the membrane with sterile water, and the cell culture inserts we placed in a tissue culture plate to establish two compartments. Each compartment was filled with RGC media and allowed to equilibrate. The upper compartment was seeded with 1 × 10^5^ hRGCs at the same time as an equal volume of media was added to the bottom compartment to limit any pressure gradients that would drive cells through the membrane. To allow for some initial cell attachment, two hours after seeding, less than 0.5 – 2.5 μL of each chemokine was added to the bottom compartment to serve as a chemoattractant (50 ng/mL aFGF (Cell Guidance Systems, GFM48), 100 ng/mL Netrin (R&D Systems, 1109-N1-025), 500 ng/ mL SHH (Cell Guidance Systems, GFM55), 1000 ng/mL Slit (R&D Systems, 5199-SL-050), and 10, 50, 200, and 500 ng/mL SDF1 (Cell Guidance Systems, GFH46). After 24 hours at standard culture conditions, the RGCs were fixed with 4% paraformaldehyde and imaged imminently on an Olympus IX83-FV3000 confocal microscope. Due to the tdTomato reporter, hRGCs could be visualized on the top and bottom of each membrane through a series of z-stacked images without needing staining. RGC recruitment was reported as the ratio of RGCs on the basal surface of the membrane to the total number of RGCs. At least three fields of view per membrane and at least four wells per group were analyzed per biological replicate. The number of cells on each side of the membrane was counted using a custom NIH ImageJ macro.

### Donor cell transplantation

All transplantation studies were approved by the Schepens Eye Research Institute of Mass. Eye and Ear Institution Animal Care and Use Committee (IACUC) following the Association for Research in Vision and Ophthalmology (ARVO) guidelines. After cell isolation, RGCs were formulated in RGC media [25 μM L-glutamic acid, 1x Glutamax, 1x Anti-Anti, 1x B27, 1x ITC-X, 5 μM forskolin in Neurobasal media supplemented with 50 ng/mL BDNF and CNTF] at 1 – 5 × 10^5^ cells/mL. The RGC suspension was then kept on ice for 1 – 2 hours to allow damaged cell membranes to seal and enable time to precondition donor cells with small molecules before transplantation. For the loss-of-function studies, RGCs were preconditioned with 10 µM AMD3100, a CXCR4 antagonist, 200 µM CK666 (Sigma, SML0006), an ST inhibitor, and/or 15 µM roscovitine (Enzo Life Sciences, ALX-380-033), an MP migration inhibitor for 1 hour on ice. Excess small molecule inhibitors were subsequently removed by centrifugation. After membrane recovery and preconditioning, donor RGCs were formulated for transplantation at 2 × 10^4^ cells/µL in RGC media containing slow-release neurotropic factors (150 U/µL GDNF-, BDNF-, and CNTF-loaded polyhedrin-based particles (PODs); Cell Guidance Systems, PPH1, PPH2, PPH59) to support cell survival in vivo.

Intravitreal (IVT) and subretinal (SR) injections were performed in young adult mice (1-to 4-month-old) under a general (ketamine/ xylazine intraperitoneal injections) and local (proparacaine eye drops) anesthetic. The donor RGC suspension (1 µL) was delivered IVT or SR through a beveled glass microneedle (80 µm inner diameter) at a 1 µL/min flow rate. Imminently following donor cell delivery, 10 ng SDF1 (1 µL) was delivered on the reverse side of the retina (SR cells – IVT SDF1 or IVT cells – SR SDF1) to establish a chemokine gradient across the neural retina. The SR space was accessed through the sclera to deliver cells or SDF1 between the retinal pigment epithelium and photoreceptor outer segments. Mice were maintained on a standard 12-hour day-night cycle and euthanized at each experimental endpoint (2 weeks for syngeneic transplantation and 3 days for xenotransplantation) by CO2 inhalation. To access the transplantation outcome and donor RGC structural integration, eyes were imminently enucleated, fixed for 24 hours at room temperature in 4% paraformaldehyde, and then processed for retinal flat-mount preparation.

### Endogenous Reprogramming

All endogenous reprogramming studies were approved by the University of Washington IACUC following the ARVO guidelines. Müller glia (MG) were reprogrammed into RGCs as previously described.^11^ Glast-CreER:LNL-tTA:tetO-mAscl1-GFP-tetO:Atoh1 adult mice were used to specifically induce Ascl1:Atoh1 in MG. Male and female mice were used in all experiments. Tamoxifen (1.5 mg per 100 mL of corn oil) was injected intraperitoneally in adult mice for 5 consecutive days to induce endogenous reprogramming. Approximately one week later, 100 µM NMDA (1.5 µL) was injected intravitreally with a 32-gauge Hamilton syringe to cause host RGC death. Two days following NMDA damage, 0, 10, or 100 ng SDF1 was injected intravitreally, and the eyes were enucleated two weeks later for analysis.

### Immunohistochemistry

To visualize host, donor, and newborn neurons in culture, retinal flat-mounts, and retinal sections, cells and tissue were fixed for 30 min for cell culture and > 4 hours for tissue samples with 4% (v/v) paraformaldehyde at room temperature. For sections with endogenous reprogramming, fixed eyes were incubated overnight in 30% sucrose at 4°C, frozen in optimal cutting temperature compound, and cryosectioned at 18 µm. For sections of wild-type retinas treated with SDF1, whole eyes were sent to the Schepens Eye Research Institute Morphology Core for paraffin embedding (6 µm sections) and Hematoxylin and eosin (H&E) staining. Paraffin-embedded sections were first deparaffinized in xylene substitute I, washed with serial dilutions of ethanol, and then demasked for 30 min using sodium citrate buffer (10 mM Sodium citrate, 0.05% tween-20, pH 6.0) before immunostaining.

For immunofluorescence staining, samples were incubated in blocking buffer [10% goat serum, 1% bovine serum albumin (BSA), 0.1% sodium citrate, 0.25% tween-20, 0.25% triton-X, and 0.3M glycine in 1x phosphate-buffered saline (PBS)] containing 1:400 goat anti-mouse overnight (>12 hours) at 4°C to limit nonspecific staining. Following blocking, RGCs and/or retinal tissue was incubated with primary antibodies in staining buffer [1% BSA, 0.25% tween-20, and 0.25% triton-X in 1x PBS] for 48 – 72 hours at 4°C. A complete list of primary antibodies and their concentrations used in this study can be found in Table S2. Each tissue sample was washed thrice with wash buffer [0.1% tween-20, 0.1% triton-X in 1x PBS] to remove any unbound primary antibody before adding the secondary antibodies. Goat anti-chicken, goat anti-rabbit, and goat anti-mouse secondaries were prepared at 1:500 in staining buffer and incubated for 12 – 24 hours at 4°C. Each sample was then again with wash buffer in triplicate and then incubated with 1 µg/mL DAPI (4’,6-diamidino-2-phenylindole) in PBS for 20 min to stain the cell nuclei. For tissue samples, tissue autofluorescence was suppressed by placing each sample in 10 mM copper sulfate (pH 5.0) for 15 min at room temperature. After a final rinse with PBS, samples were mounted between glass cover slides and #1 coverslips with a polyvinyl alcohol-based mounting media containing an antifade reagent (1,4-Diazabicyclo[2.2.2]octane). Samples containing stem cell-derived RGCs and/or wild-type retinas were imaged on an Olympus IX83-FV3000 confocal microscope. Retinas with endogenous reprogramming were imaged on a Zeiss LSM880 confocal microscope at 20x magnification.

### Flow Cytometry

To quantify the percentage of donor RGCs that express CXCR4, dissociated cells were incubated on ice in isolation buffer [2 mM EDTA, 2% FBS, and 0.1% BSA in Hank’s Balanced Salt Solution without Ca^2+^ and Mg^2+^] with 1:200 mouse anti-CXCR4 (BL304502). After a 20 min incubation on ice, cells were centrifuged at 200 x g for 5 min and resuspended in isolation buffer to remove any unbound primary antibody. Following the resuspension of the cell pellet, 1:400 goat anti-mouse AlexaFluor488 (AF488) was added to the cell suspension for 10 minutes on ice to visualize the CXCR4 receptor. To remove all the unbound secondary antibodies from the cell suspension, the RGCs were centrifuged at 200 x g for 5 min and resuspended in isolation buffer. After this wash, the cell suspension was filtered through a 40 µm cell strainer, and flow cytometry was performed on a BD LSR II analyzer. The data were gated using the FlowJo software for single-cells using forward/side scatter. Lastly, gates were drawn to identify the donor RGCs (Brn3b-tdTomato) and CXCR4 (AF488) expressing cells to calculate the percentage of CXCR4 positive RGCs.

### Assessment of RGC distribution

Donor RGC position within the host neural retina was quantified with custom semi-automated ImageJ and MATLAB scripts. Retinal flat-mounts were imaged using the resonate scan on an Olympus IX83-FV3000 confocal microscope at 20x magnification (UPlanSApo). Z-axis intervals were set to 2 µm, and mosaic tiled images of the whole retinal flat-mount were taken at a 512×512 pixel resolution per tile. To identify donor RGCs within the host retina, each tile from the retinal flat-mount was first imported into ImageJ as an image sequence. After subtracting the background, the 3D object counter plugin was then used to export the size and local position of donor RGCs in each tile to a ‘.csv’ file. The MATL file containing the map data with the positions of each title relative to each other was manually saved as a ‘.txt’ file for every retina. These files were imported into MATLAB to calculate the global x, y, and z-positions of each identified donor RGC in addition to the total number of donor RGCs per retina. These coordinates were used to generate depth-coded color maps that could be manually overlayed on their corresponding retinal flat-mount. The number of cells within a cell cluster was calculated by dividing the volume of that cluster by 1.5x the average donor RGC volume. To calculate the percentage of donor RGC that migrated into the GCL, the z-axis position of the GCL was manually inputted into our MATLAB script. The GCL’s position was identified by randomly selecting four tiles from each retinal flat-mount and averaging the z-plane containing RBMPS+ host cells. To compare z-positions of donor RGCs across retinas, the z-position was normalized between 0 and 100 using min/max scaling. Finally, donor RGC coverage was calculated as the percentage of tiles per retina with one or more donor RGC.

### Quantification of in situ displacement

The positions of MG-derived neurons were identified by the co-expression of GFP (pseudo-colored white) and HuC/D (red) using a custom ImageJ macro. Each image from the retinal section was first imported into ImageJ as an image sequence, then processed using the Auto Threshold function. To find the coordinates of HuC/D positive MG, binary masks were created using the red channel and multiplied to the white channel so that only MG with HuC/D expression were visible. This image was then processed using the Auto Threshold and Analyze Particles functions to export the coordinates of the HuC/D positive MG. The normalized displacement error (NDE) was then calculated from each image in three steps: First, sorting the identified cells according to their x-axis positions and summing the total length between them; second, subtracting this segment length that connects each cell by the straight line that connects the first and last cell in series; and third, normalizing this difference by dividing by the total number of cells. At least three images per retina were analyzed to calculate the NDE.

### Statistical Analysis

Statistical significance was calculated using GraphPad PRISM 9 using a Tukey one-way ANOVA. Error bars represent the mean ± standard deviation (SD) of measurements (*p < 0.05, **p < 0.01, ***p < 0.001, ****p < 0.0001). Each dot within a bar plot represents an individual retina or well, while each dot with a violin plot represents an individual cell.

### Data availability

The raw single-cell RNA sequencing data of human fetal retina and organoids from Sridhar et al. (2020) and human adult retina from Wang et al. (2022) used in this study are available in the GEO database under accession code GSE142526 and GSE196235. The processed RNA sequencing data generated in this study are available on GitHub at: https://github.com/mcrewcow/BaranovLab

## Funding

NIH/NEI – 1F32EY033211 (JRS)

NIH/NEI – 5T32EY007145 (MBED fellowship, JRS)

NIH/NEI K99EY033402 (LT)

NIH/NEI – 5U24EY029893-03 (PB)

NIH/NEI – P30EY003790 (Core Facility Grant)

Bright Focus Foundation – G2020231 (PB)

NIH/NEI R01EY021482 (TAR)

Gilbert Family Foundation – GFF00 (PB & TAR)

## Acknowledgements

The authors would like to thank Connor Finkbeiner for their advice in processing the raw single-cell RNA sequencing fetal and organoid data, Joshua Sanes for Thy1-GFP mice, Don Zack for providing the Brn3b-tdTomato hESCs, and Randy Huang for flow cytometry support.

## Author contributions

JRS and PB conceived the project and designed experiments. JRS performed the in vitro assays, immunohistochemistry, confocal microscopy, and quantitative transplant analysis. JRS and MP differentiated RGCs. PB performed the cell transplantation. LT performed the endogenous reprogramming experiment under the advisement of TAR. EK analyzed the single-cell RNA sequencing data. JRS wrote the manuscript and prepared the figures. All authors edited and provided feedback on the manuscript. PB supervised the work.

## Competing interests

The University of Washington has a patent incorporating the endogenous reprogramming technology described in this report with inventors LT and TAR.

## Supplemental Information

Fig. S1-S12

Table S1-S2

**Fig. S1.**
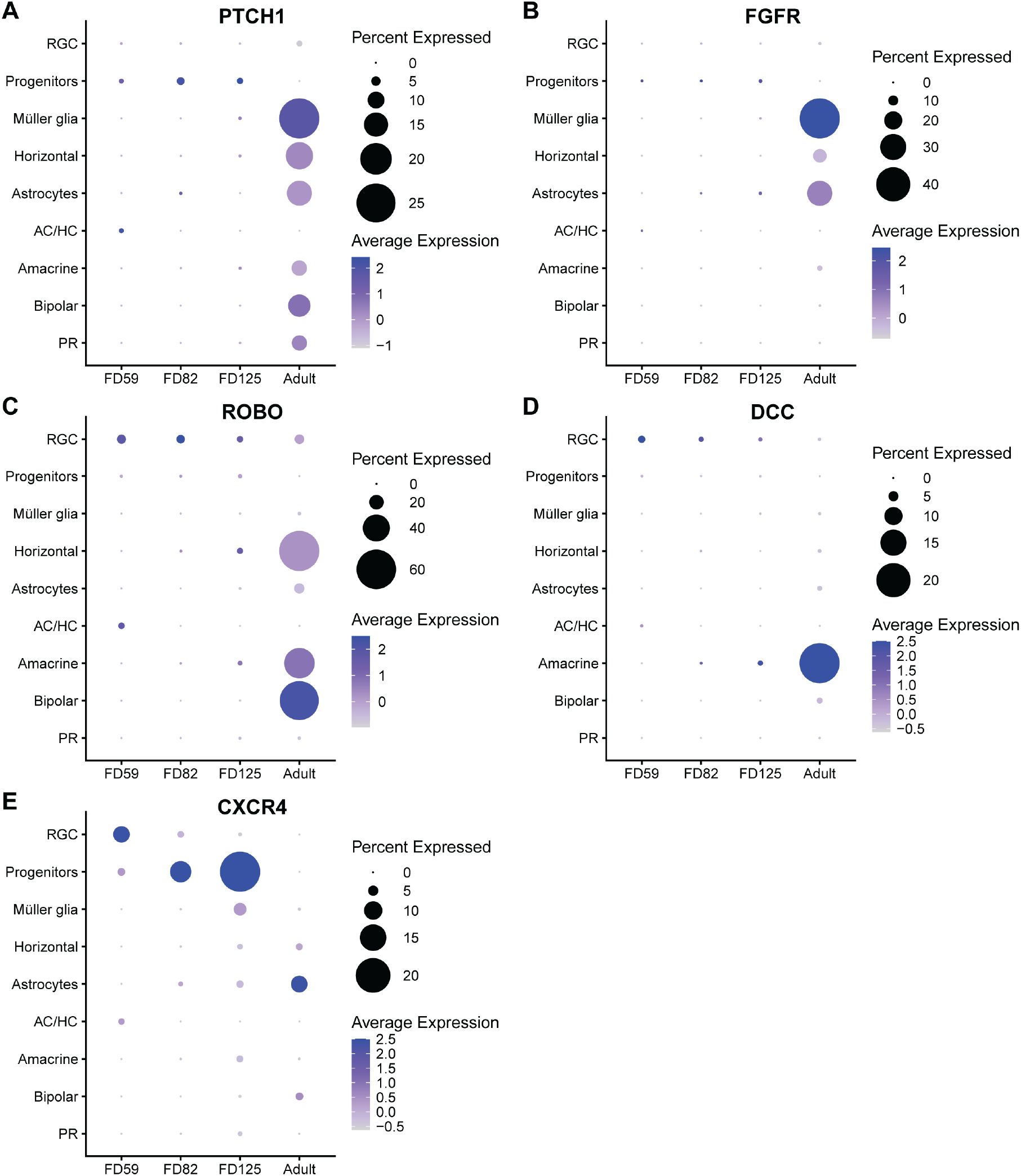
Human retinal neuron chemokine expression profiles during development. **(A)** Dot plots showing the percent and average expression of PTCH1, **(B)** FGFR, **(C)** ROBO, **(D)** DCC, and **(E)** CXCR4 in the human retina at different stages of development (fetal days 59, 82, and 125, and adult) for each neural population.

**Fig. S2.**
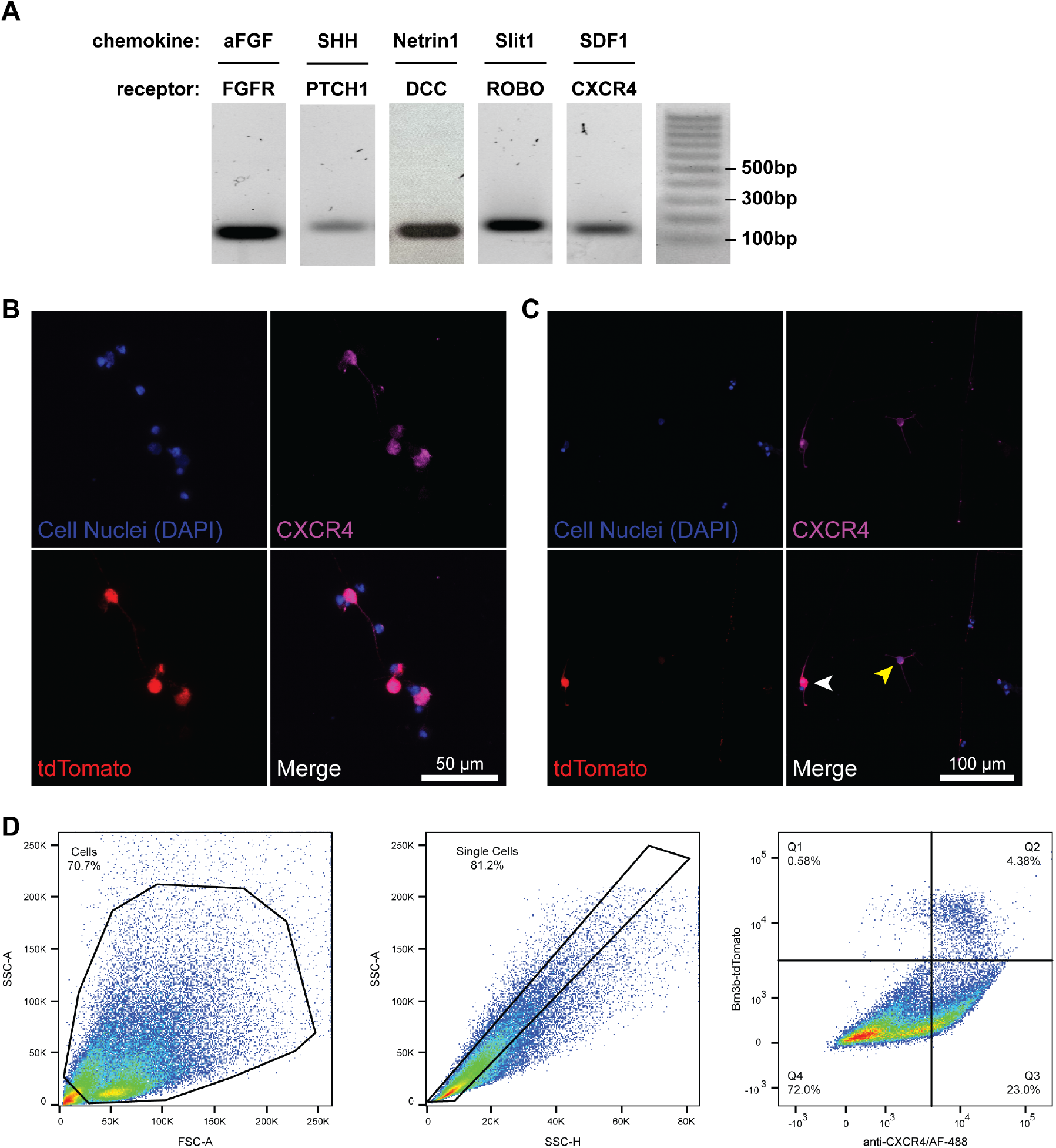
Stem cell-derived RGCs express chemokine receptors. **(A)** Images of reverse transcription PCR products from sorted stem cell-derived RGCs show the expression for FGFR, PTCH1, DCC, ROBO1, and CXCR4. **(B)** Representative immunofluorescent images of cultured stem cell-derived RGCs show all tdTomato positive RGCs (red) co-express CXCR4 (violet), but **(C)** not all CXCR4 positive cells are tdTomato positive (yellow arrow: CXCR4 positive, tdTomato negative; white arrow: CXCR4 positive, tdTomato positive). Cell nuclei are stained for DAPI (blue). **(D)** FACS plots with gating confirms that > 85% of RGCs (tdTomato positive) express CXCR4 (AF488 positive).

**Fig. S3.**
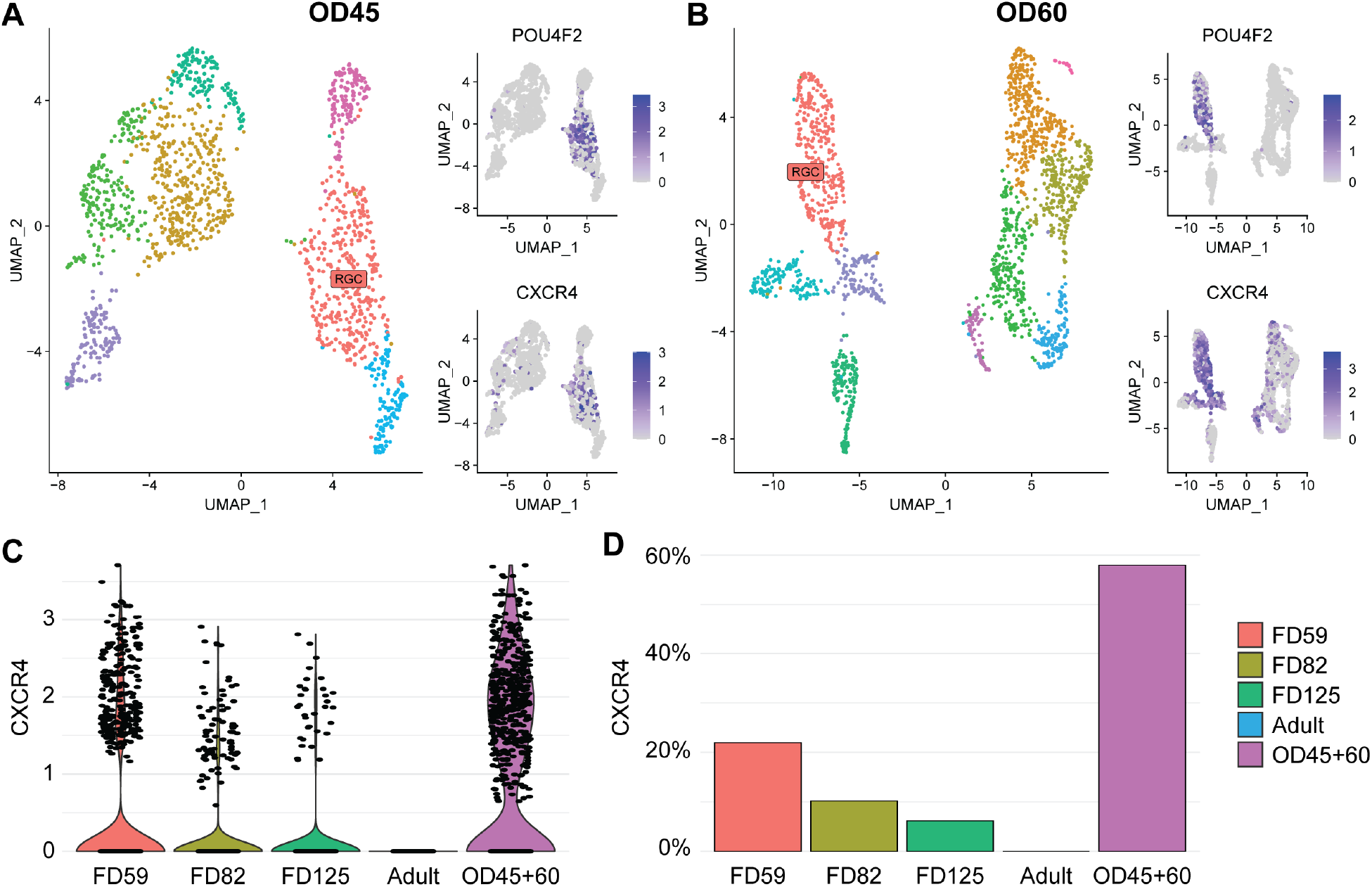
Characterization of human RGC CXCR4 expression during in vivo and in vitro development. **(A)** UMAP plot of human retinal organoids with RGC clusters identified by cell-type-specific gene expression and feature plots showing characteristic genes describing the cluster of RGCs (Pou4f2) and CXCR4 after 45 days **(B)** and 60 days of organoid differentiation. **(C)** Single-cell quantification and **(D)** percentage of CXCR4 expression in RGCs during human development compared to retinal organoid differentiation shows increased expression in organoids compared to human tissue.

**Fig. S4.**
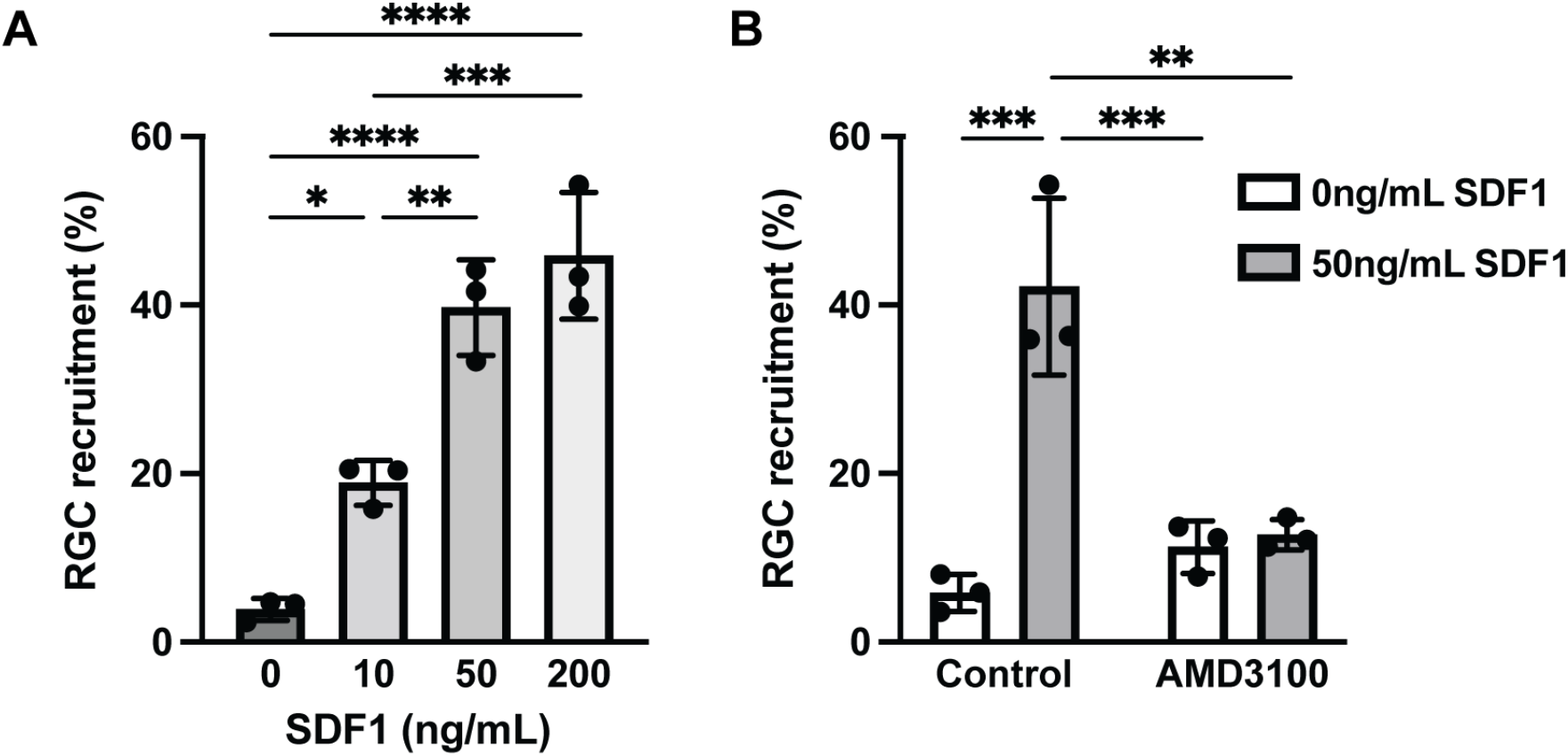
SDF1-mediated RGC recruitment is dose-dependent and functions through CXCR4. **(A)** Quantitative analysis of RGC recruitment, defined as the ratio of RGCs on the basal surface to the total number of RGCs, in response 0, 10, 50, and 200 ng/mL SDF1, and/or **(B)** AMD3100 (CXCR4 antagonist) treatment. N = 3 wells per group.

**Fig. S5.**
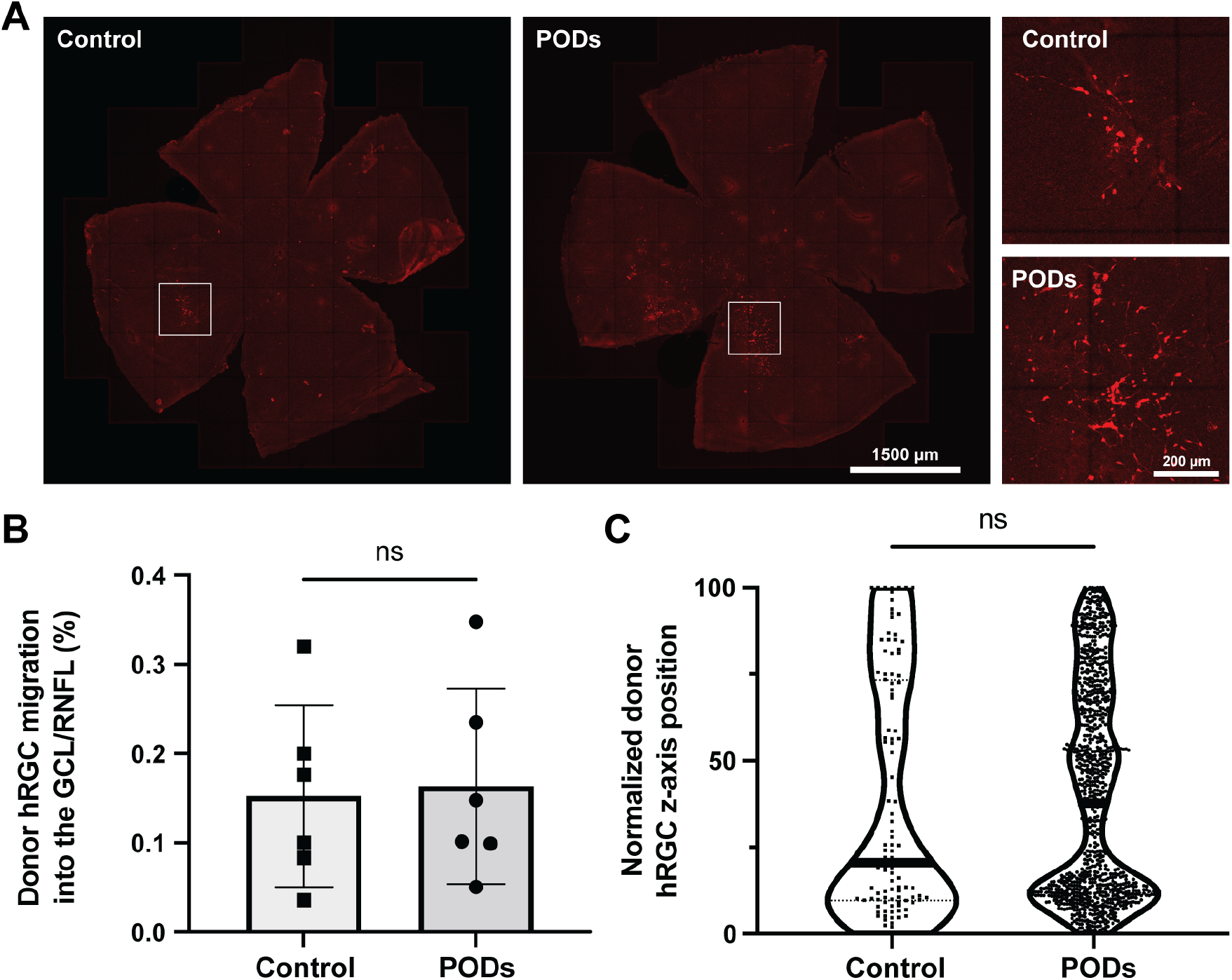
Human stem cell-derived RGC xenotransplantation survival is improved by slow-release neurotropic factors. **(A)** Representative retinal flat mount (max intensity projection) showing donor RGCs within the neural retina following transplantation with or without the inclusion of slow-release neurotropic factors (GDNF-, BDNF-, and CNTF-loaded polyhedrin-based particles (PODs)) in the cell delivery formulation. Inlay (zoomed-in region) shows RGC morphology and increased neurite outgrowth in the PODs group. **(B)** Quantitative analysis of donor RGC migration into the GCL/RNFL shows no significant increase in donor RGCs translocating into the GCL/RNFL with PODs formulation. N = 6 mice per group. **(C)** Single-cell quantification of z-axis position normalized to the thickness of the retina shows that the inclusion of PODs within the cell delivery formulation does not affect donor RGC migration.

**Fig. S6.**
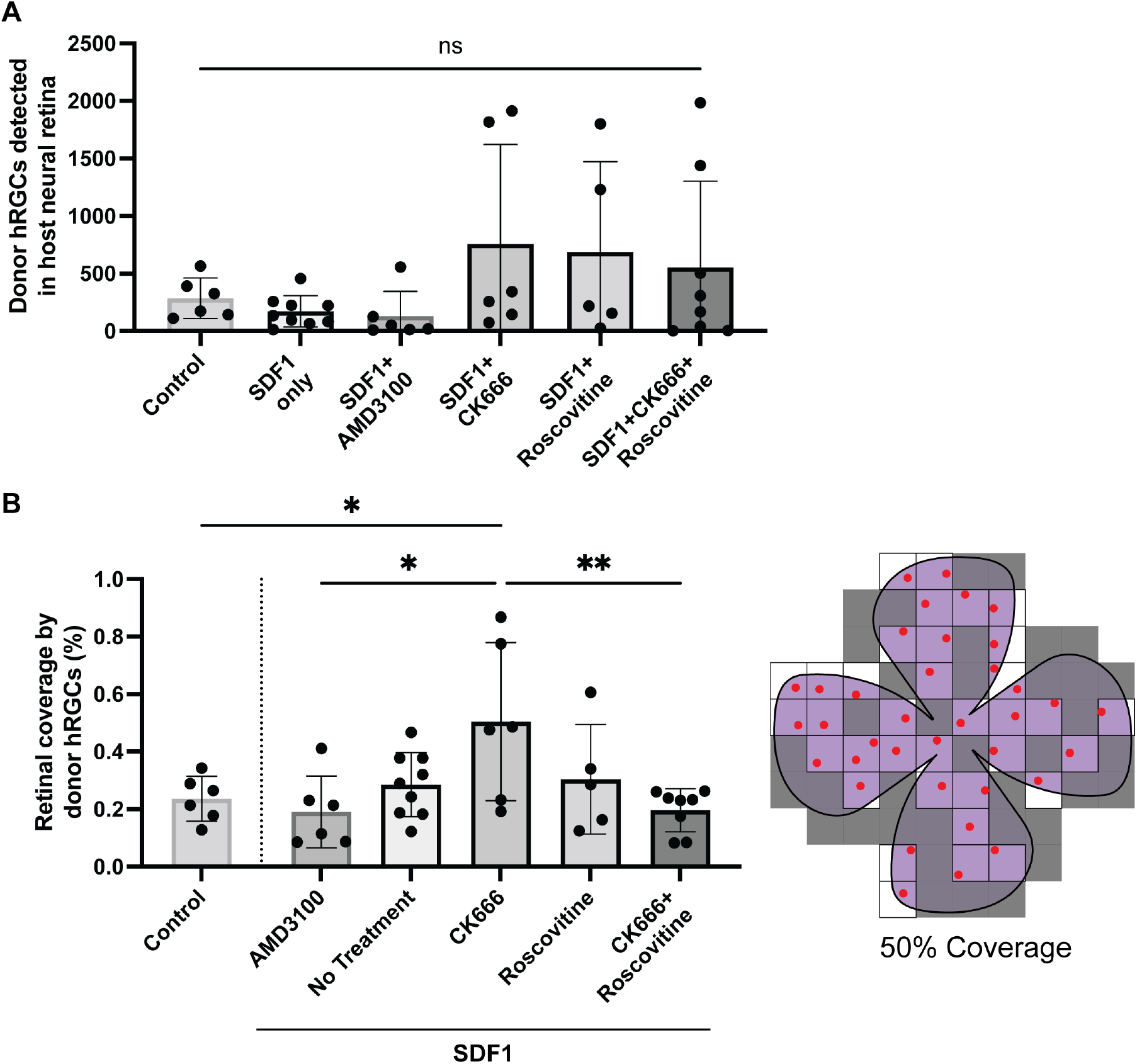
Donor RGC numbers and coverage within the neural retina. **(A)** The number of donor RGCs detected in the neural retina shows no statistically significant differences between treatments and donor RGC preconditioning. **(B)** Quantitative analysis of donor RGC coverage, defined as the percentage of tiles per retina that have one or more donor RGC (i.e., 50% coverage representation), shows an increase in coverage for donor RGCs preconditioned with CK666, an ST inhibitor.

**Fig. S7.**
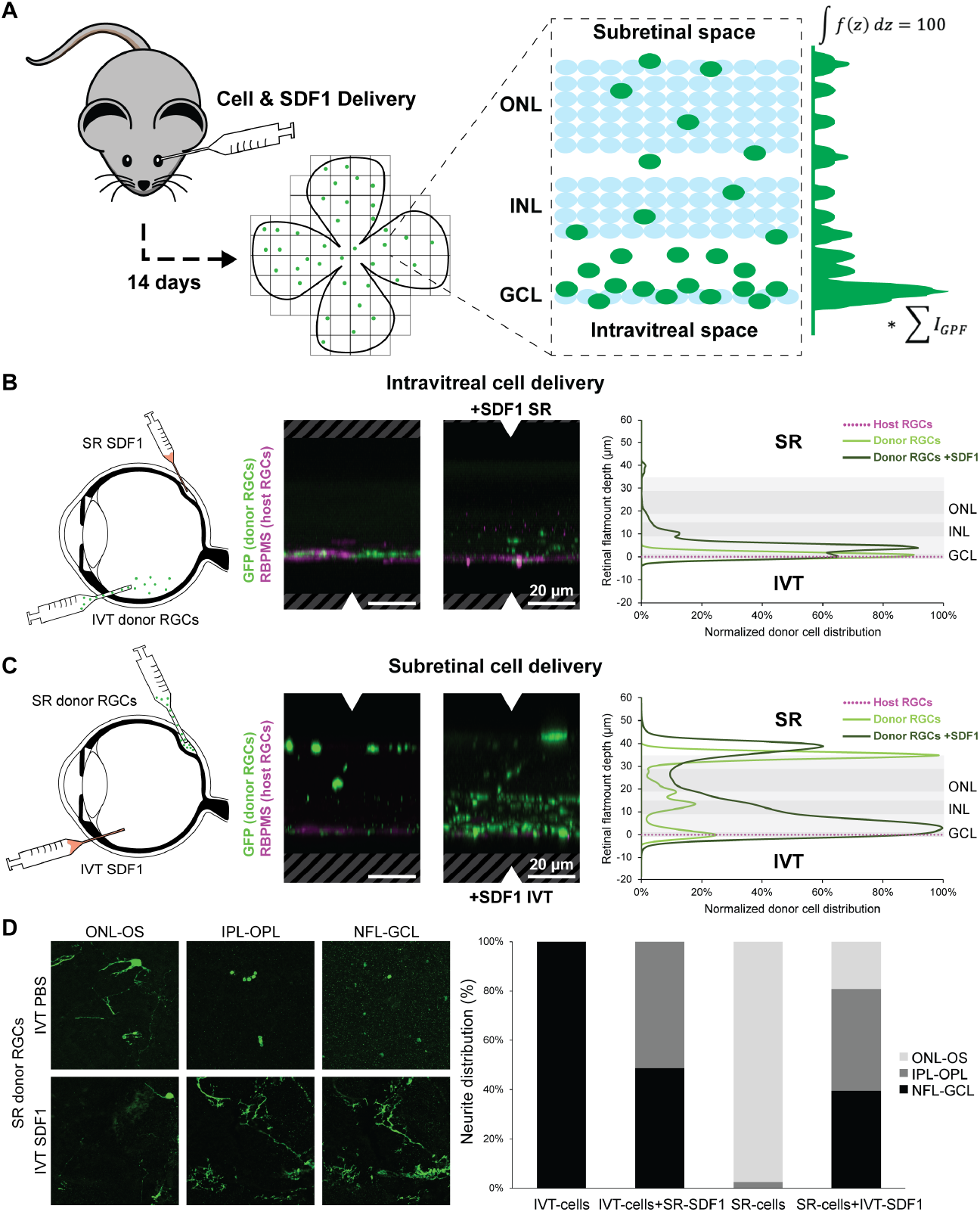
Mouse stem cell-derived syngeneic transplantation in response to SDF1 treatments. **(A)** Schematic overview of our transplantation and quantification strategy to assess the donor cell distribution 14 days following transplantation using orthographic projections of a retinal flat mount. Donor cell distribution was calculated by setting the area under the intensity profile to 100% and multiplying by the normalized (total/maximum) GFP intensity. **(B)** Schematic representation of delivery strategy to establish a chemokine gradient across the neuronal retina following IVT RGC and **(C)** SR RGC delivery. Representative immunofluorescent images of retinal flat mount orthographic projections stained for the donor (GFP, green) and host RGCs (RBPMS, violet) and quantifying normalized donor cell distribution show donor RGC migration/neurite extension in the direction SDF1 irrespective of the direction of the gradient. **(D)** Representative immunofluorescent images showing the neurites of donor RGCs (GFP, green) within different layers of the neural retina in response to SDF1 following SR delivery. Quantifying neurite distribution within each retinal layer between groups shows that SDF1 treatment results in increased neurite innervation within the retina. ONL: outer nuclear layer; OS: outer segments; OPL: outer plexiform layer; INL: inner nuclear layer; inner plexiform layer; GCL: ganglion cell layer; NFL: nerve fiber layer.

**Fig. S8.**
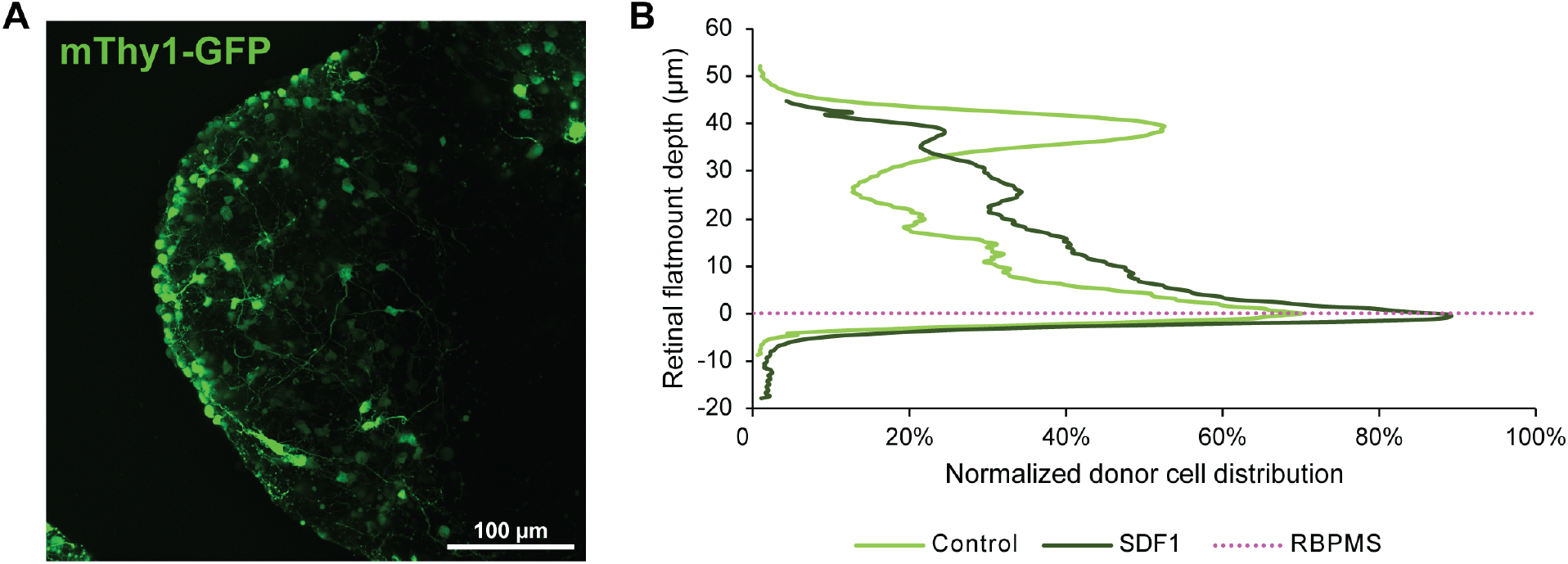
Whole retinal organoid transplantation in mice with SDF1 treatments. **(A)** Representative live-cell image of mouse retinal organoids containing RGCs (recombinant GFP expression, green). **(B)** Quantifying normalized donor cell distribution shows donor RGC migration/neurite extension out from the retinal organoids in the SR space in the direction of IVT SDF1. Normalized donor cell distribution was calculated by setting the area under the intensity profile to 100% and multiplying by the normalized (total/maximum) GFP intensity.

**Fig. S9.**
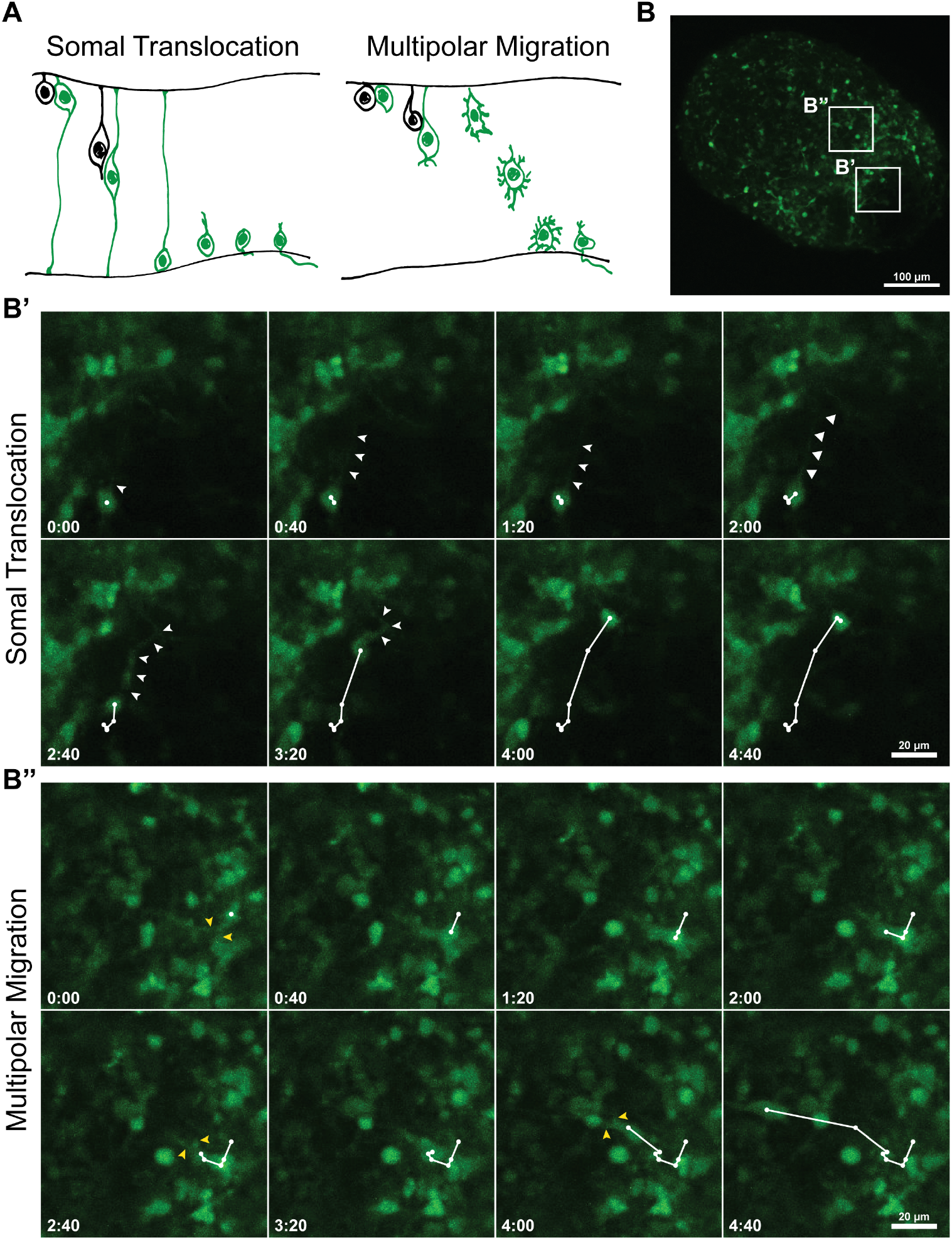
RGC migrate within 3D stem cell-derived retinal organoids by two independent patterns of neuronal translocation. **(A)** Graphical illustration of RGC ST and MP migration towards the basal surface. **(B)** Representative live-cell images of a 3D mouse retinal organoid (max intensity projection) containing differentiated RGCs are indicated by the recombinant GFP expression under a sparse Thy1.2 promoter. **(B’)** The zoomed-in region shows the ST and **(B’’)** MP migration patterns within a retinal organoid. The white arrows highlight the unipolar extending neural process in ST, and the yellow arrows point to the multipolar neurite extension in MP migration. The trajectories are recorded at 20 min intervals.

**Fig. S10.**
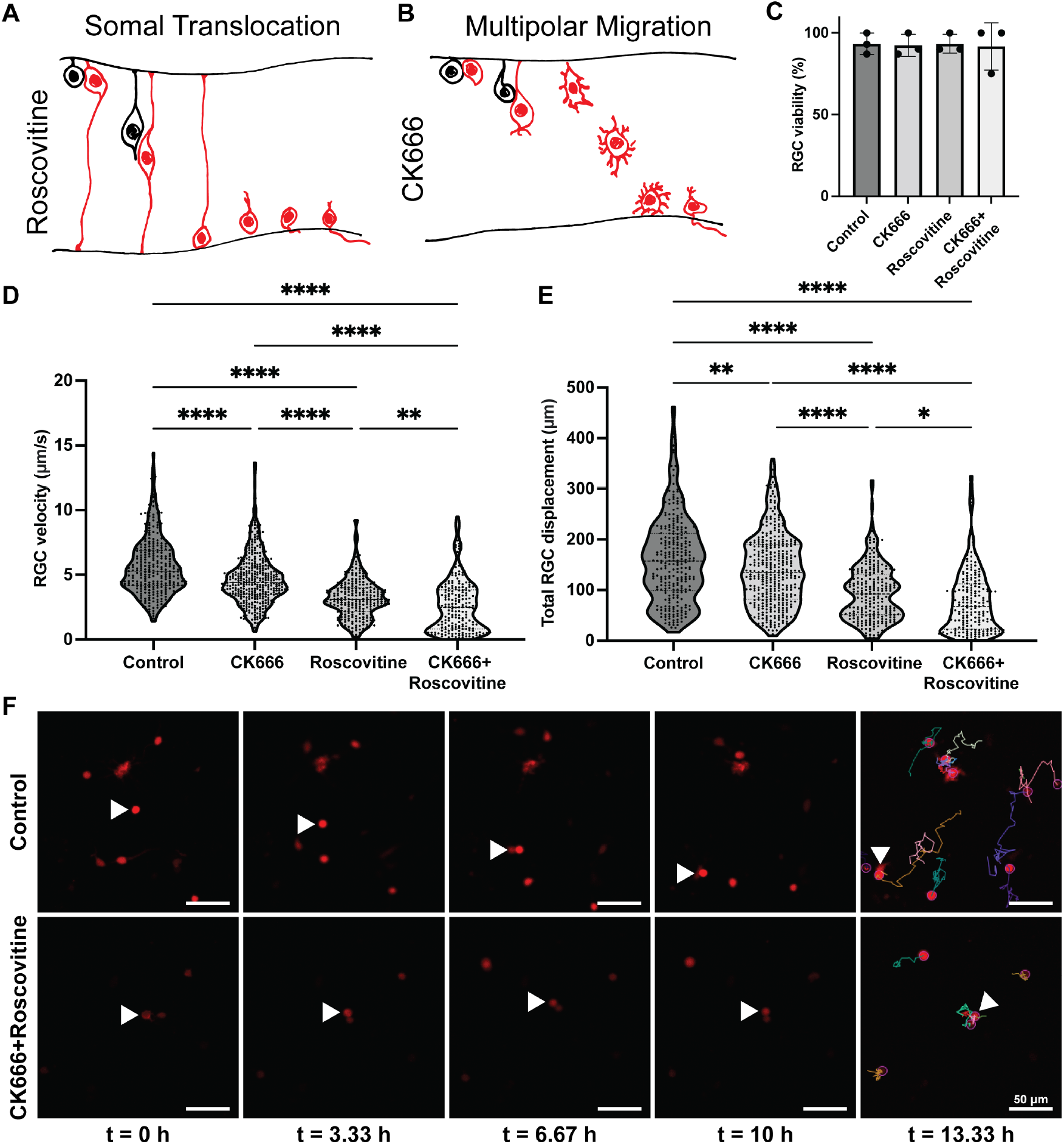
Human stem cell-derived RGC migrate by two independent modes of migration in vitro. **(A)** Graphical illustration of RGC ST towards the basal surface resulting from roscovitine treatment, an MP migration inhibitor. **(B)** Graphical illustration of RGC MP migration towards the basal surface resulting from CK666 treatment, an ST inhibitor. **(C)** Quantifying cell viability by calcein-AM/ethidium homodimer staining demonstrates that small molecule migration inhibitors do not affect RGC viability. **(D)** Violin plots of RGC velocity and **(E)** total displacement for RGCs tracked on laminin-coated plates in response to each inhibitor molecule show distinct migration profiles between groups with a near complete inhibition of migration when RGCs are treated with both CK666 and roscovitine. **(F)** Representative live-cell time-lapse images show the trajectories of RGCs (recombinant tdTomato expression, red) with high (control) and low (CK666 + roscovitine) migration activity.

**Fig. S11.**
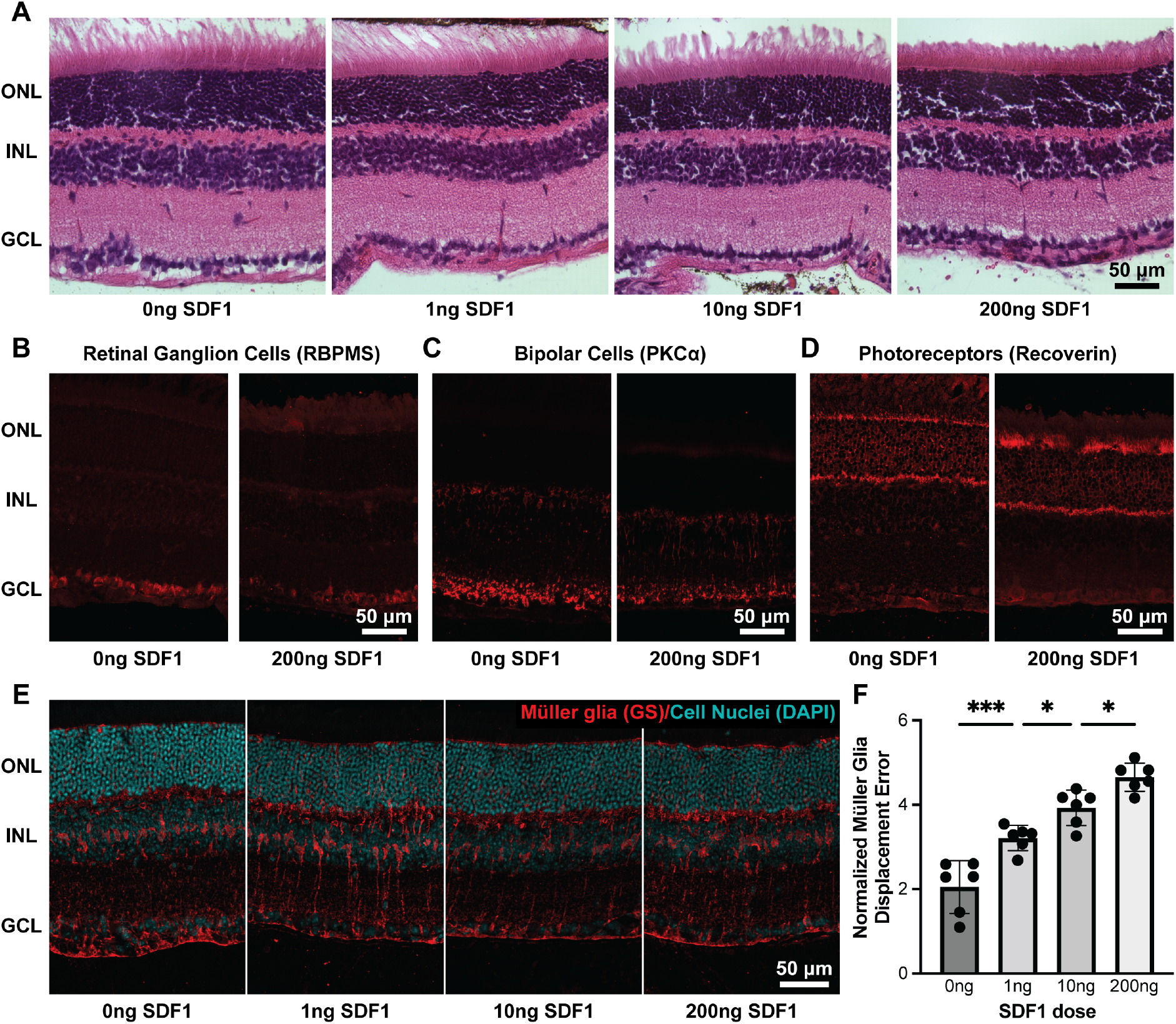
The effect of SDF1 on retinal lamination and position of host retinal neurons. **(A)** Representative hematoxylin and eosin (H&E) stained sections of the neural retina following intravitreal injection of 0, 1, 10, and 200 ng SDF1 show SDF1 does not alter retinal lamination. **(B)** Representative immunofluorescent images of retinal sections stained for host RGCs (RBPMS positive, red), **(C)** bipolar cells (PkCa positive, red), and **(D)** and photoreceptors (recoverin positive, red) following intravitreal injection of 0 and 200 ng SDF1 shows no effect of SDF1 on the position of host retinal neurons. **(E)** Representative immunofluorescent images of retinal sections stained for host Müller glia (GS positive, red) and cell nuclei (DAPI, cyan) show that SDF1 affects the displacement of Müller glia in the INL. (F) Quantitative analysis of normalized Müller glia displacement error in response to intravitreal injection of 0, 1, 10, and 200 ng SDF1 shows increased displacement with increasing concentrations of SDF1. Normalized Müller glia displacement error is calculated as the difference between the total segment length that connects all the cell bodies and a straight line normalized to the total number of cells. ONL: outer nuclear layer; INL: inner nuclear layer; GCL: ganglion cell layer.

**Fig. S12.**
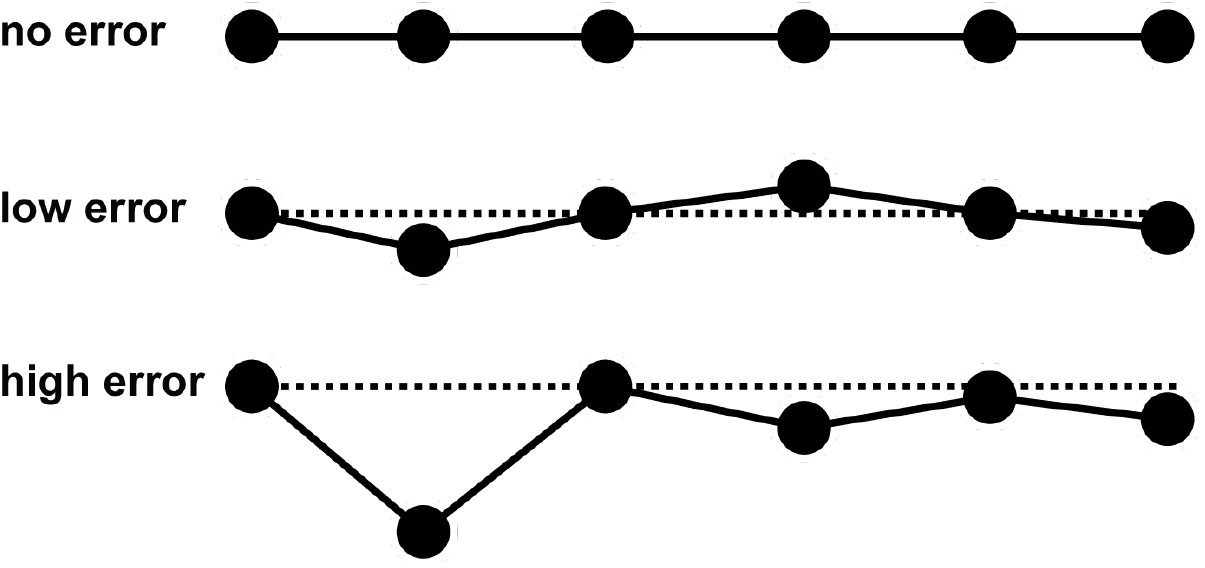
Displacement error calculation. Schematic representation of no, low, and high displacement error, defined as the difference between the total segment length that connects all the cell bodies and a straight line.

**Table S1.** Software packages for single-cell RNA sequencing analysis.

| Package | Version | Link |
| --- | --- | --- |
| biomaRt | 2.48.3 | https://bioconductor.org/packages/release/bioc/html/biomaRt.html |
| cowplot | 1.1.1 | https://cran.r-project.org/web/packages/cowplot/ |
| dittoSeq | 1.4.4 | https://bioconductor.org/packages/release/bioc/html/dittoSeq.html |
| dplyr | 1.08 | https://dplyr.tidyverse.org/ |
| EnhancedVolcano | 1.10.0 | https://bioconductor.org/packages/release/bioc/html/EnhancedVolcano.html |
| escape | 1.2.0 | https://github.com/ncborcherding/escape |
| ggplot2 | 3.3.5 | https://ggplot2.tidyverse.org/ |
| GSEABase | 1.54.0 | https://bioconductor.org/packages/release/bioc/html/GSEABase.html |
| htmlwidgets | 1.5.4 | https://www.htmlwidgets.org/ |
| Matrix | 1.3-4 | https://cran.r-project.org/web/packages/Matrix/index.html |
| patchwork | 1.1.1 | https://patchwork.data-imaginist.com/ |
| plotly | 4.10.0 | https://plotly.com/r/ |
| reticulate | 1.24 | https://rstudio.github.io/reticulate/ |
| Seurat | 4.1.0 | https://satijalab.org/seurat/ |
| SeuratDisk | 0.0.0.9019 | https://github.com/mojaveazure/seurat-disk |
| SeuratWrappers | 0.3.0 | https://github.com/satijalab/seurat-wrappers |

**Table S2.** Primary antibodies for immunohistochemistry.

| **Antibody** | **Concentration** | **Vendor** | **Catalog #** |
| --- | --- | --- | --- |
| Chicken Anti-mCherry | 1:500 | Abcam | ab205402 |
| Chicken Anti-GFP | 1:50 | enQuire BioReagents | NC1666473 |
| Rabbit Anti-RBPMS | 1:300 | GeneTex | GTX118619 |
| Chicken Anti-GFP | 1:2000 | Abcam | ab13970 |
| Rabbit Anti-CXCR4 | 1:250 | Abcam | ab181020 |
| Mouse Anti-PKCa | 1:100 | Santa Cruz | sc8392 |
| Rabbit Anti-Recoverin | 1:100 | Millipore | AB5585 |
| Rabbit Anti-GS | 1:200 | Abcam | ab49873 |
| Mouse Anti-HuC/D | 1:250 | Invitrogen | A21271 |

